# Evolutionarily Conserved Decline of tRNA Mannosyl-Queuosine Links Translational Regulation to Aging and Is Reversed by Queuine

**DOI:** 10.64898/2026.03.22.713446

**Authors:** Rui-Ze Gong, Tong-Meng Yan, Yu Pan, Kai-Yue Cao, Ya-Ting Cheng, Liu-Ying Mo, Zhi-Hong Jiang

**Affiliations:** State Key Laboratory of Mechanism and Quality of Chinese Medicine, Macau University of Science and Technology, Macau 999078, China

**Keywords:** Aging hallmark, tRNA modification, manQ, queuine, anti-aging

## Abstract

Aging arises from interconnected molecular defects, yet upstream regulatory mechanisms that coordinate these hallmarks remain incompletely defined. While epitranscriptomic regulation has emerged as a critical layer of gene control, the contribution of tRNA-specific modifications to aging remains largely unexplored. Here, we systematically profile tRNA modifications across multiple organs, species, and senescence models and identify mannosyl-queuosine (manQ), a wobble-position modification of tRNA^Asp^, as the first tRNA-specific modification that consistently declines with age. ManQ depletion is evolutionarily conserved and tightly correlates with functional deterioration. Mechanistically, loss of manQ impairs translational fidelity, leading to proteome imbalance, collapse of proteostasis, and aberrant expression of senescence-associated proteins, including GPNMB. These translational defects intersect with established aging hallmarks and accelerate cellular and organismal aging. We further demonstrate that circulating queuine, a microbiota-derived precursor required for manQ biosynthesis, declines with age in rodents and humans. Queuine deficiency promotes senescence, whereas supplementation restores manQ levels, improves translational accuracy, suppresses p16/p21-driven senescence programs, and re-establishes proteostatic balance. Across species, queuine supplementation extends lifespan and enhances healthspan. In Drosophila, it increases median lifespan by 47% and improves stress resistance and memory. In naturally aging mice, long-term oral administration extends lifespan by 15.3%, reduces DNA methylation age, improves cognitive and motor performance, strengthens antioxidant defenses, remodels the gut microbiota, and alleviates inflammation and metabolic dysfunction without detectable toxicity. Collectively, these findings establish tRNA epitranscriptomic remodeling as a previously unrecognized layer of aging regulation and identify restoration of manQ through queuine supplementation as a multi-system strategy to delay aging.

## INTRODUCTION

Aging constitutes a complex physiological process characterized by progressive functional decline, diminished adaptive capacity across molecular, cellular, tissue, and organ systems, and an elevated mortality risk (Gladyshev et al., 2021; López-Otín et al., 2023). Demographic projections estimate that by 2050, the global population aged over 60 years will reach 2.1 billion, constituting 22% of the total population. This unprecedented demographic shift poses escalating health and socioeconomic challenges (Navaneetham and Arunachalam, 2023). Consequently, aging research represents not only a frontier in fundamental biology but also a strategic priority for societal resilience. Healthspan extension has thus emerged as both a perennial human aspiration and a central focus of contemporary geroscience. Moreover, systematic elucidation of the molecular networks governing aging is crucial to deciphering senescence regulation and developing targeted geroprotective strategies (López-Otín et al., 2013). This paradigm transition, from reactive disease treatment to proactive aging modulation, holds transformative potential for achieving a pivotal biomedical milestone: realizing humanity’s shared vision of “aging without frailty”.

The landmark 2023 review in Cell systematically defined twelve conserved hallmarks of aging, spanning genomic instability to mitochondrial dysfunction, recently expanded to include two emerging features: extracellular matrix alterations and psychosocial isolation (Kroemer et al., 2025; López-Otín et al., 2023). Collectively, these hallmarks constitute an intricately interconnected network that profoundly informs our understanding of aging etiology and intervention strategies (Bao et al., 2023). However, critical gaps persist in their mechanistic interpretation and translational application. While these classical hallmarks have catalyzed the development of aging assessment frameworks, their utility in deciphering real-time regulatory dynamics remains limited, and identifying actionable targets for therapeutic intervention remains challenging (Bao et al., 2023). Novel geroprotective strategies, including caloric restriction, fecal microbiota transplantation (FMT), and heterochronic parabiosis, have garnered significant attention for their lifespan-extending potential (Boehme et al., 2021; Ma et al., 2022; Solon-Biet et al., 2020). However, prohibitive medical costs and inconsistent therapeutic efficacy severely hinder their clinical translation. In parallel, mechanistic insights into pharmacological interventions such as rapamycin, metformin, and spermidine continue to advance. Rapamycin extends healthspan in model organisms primarily through mTORC1 inhibition (Mannick et al., 2023). Spermidine delays cellular senescence by inducing autophagy and potentiating antioxidant responses (Zhang et al., 2023). Quercetin, a recognized senolytic agent, selectively eliminates senescent cells (Novais et al., 2021). Despite promising preclinical results, the clinical application of these compounds faces unresolved controversies regarding human efficacy and safety. Rapamycin’s therapeutic utility is constrained by dose-dependent immunosuppressive effects and metabolic complications (Mannick et al., 2023). Although efficacious in cellular and animal models, spermidine and quercetin lack robust validation in large-scale human clinical trials (López-Otín et al., 2023). Critically, single-target approaches inadequately address the multifactorial pathogenesis of aging, underscoring the need for combinatorial strategies that concurrently target multiple hallmarks of senescence.

Epitranscriptomics investigates post-transcriptional gene regulation through dynamic RNA modifications, offering unique insights into aging biology (McMahon et al., 2021; Wu et al., 2023). The field gained significant momentum when He et al. identified FTO as the first m^6^A RNA demethylase, establishing RNA modifications as a reversible regulatory layer (Jia et al., 2011). Subsequent research has elucidated the dynamic regulatory roles of diverse RNA modifications in disease pathogenesis and aging progression (Roundtree et al., 2017). For instance, METTL3 deficiency in human pluripotent stem cells triggers premature senescence and apoptosis (Wu et al., 2023). Conversely, TRDMT1, the m^5^C methyltransferase, alleviates cellular senescence by orchestrating DNA damage repair through recruitment of RAD51/RAD52 complexes (Chen et al., 2020). Despite these advances, current research disproportionately focuses on mRNA modifications, while the epitranscriptomic regulation of aging by tRNA modifications remains largely unexplored. tRNAs contain the most diverse repertoire of RNA modifications, with over 70% of known epitranscriptomic modifications localized to these molecules (Lorenz et al., 2017). These modifications are crucial for maintaining tRNA structural stability and ensuring translational fidelity (Pan, 2018). Accumulating evidence demonstrates that dysregulated tRNA modifications are linked to age-related pathologies, particularly neurodegenerative disorders, highlighting their systemic importance in aging (Jonkhout et al., 2017; Zhou et al., 2021). Recent studies have identified that METTL1, the methyltransferase responsible for tRNA m^7^G modification, exhibits age-dependent depletion (Fu et al., 2024). This deficiency leads to tRNA destabilization, ribosomal stalling at specific codons, and activation of ribotoxicity and integrated stress responses, ultimately driving exacerbated senescence-associated secretory phenotypes (SASP). While these findings establish tRNA epitranscriptomics as a critical regulator of aging, the field remains nascent. Notably, the mechanistic roles of tRNA-specific modifications, particularly those distinct from modifications shared with mRNA, within aging pathways have not yet been systematically characterized.

Mannosyl-queuosine (manQ), first characterized in the 1970s, is a hypermodified nucleoside uniquely found at position 34 of the anticodon loop in tRNA^Asp(manQUC)^ of mammals (Hillmeier et al., 2021; Okada and Nishimura, 1977). Despite its early discovery, both its structural conformation and biosynthetic pathway remained unresolved until recent advances (Hillmeier et al., 2021). The identification of GTDC1 as the enzyme responsible for manQ biosynthesis marked a pivotal breakthrough, thereby addressing a critical knowledge gap that had impeded mechanistic studies (Zhao et al., 2023). Strikingly conserved evolutionary patterns emerge in manQ’s developmental regulation: Thomas et al. reported significantly higher manQ levels in adult mouse kidneys and other organs compared to neonates, with similar patterns observed in rat liver models (Costa et al., 2004; Thumbs et al., 2020). Zebrafish *GTDC1* knockout studies further demonstrated severe growth defects, establishing manQ as indispensable for vertebrate post-embryonic development (Zhao et al., 2023). Queuine, the 7-deazaguanine-derived essential precursor of manQ, presents a unique nutritional paradox: while exclusively produced by gut microbiota, it is required for the maturation of both mitochondrial and cytoplasmic tRNA^Asn^, tRNA^Asp^, tRNA^His^, and tRNA^Tyr^ in mammals (**Fig. S1A, B**) (Fergus et al., 2015). Enzymatic specialization governs queuosine modification: GTDC1 catalyzes manQ formation at tRNA^Asp^ position 34, whereas B3GNTL1 generates galactosyl-queuosine (galQ) at tRNA^Tyr^ position 34 (**Fig. S1C**) (Zhao et al., 2023). Beyond conferring ribonuclease resistance to tRNAs, queuine enhances translational accuracy through codon-anticodon optimization (Kulkarni et al., 2021; Tuorto et al., 2018; Wang et al., 2018). Clinical correlations link queuine deficiency to ulcerative colitis and cognitive impairment, and Ames designate it as a “longevity vitamin” (Ames, 2018; Cirzi et al., 2023; Zhang et al., 2023). Nevertheless, its potential anti-aging properties and underlying molecular mechanisms remain uncharacterized.

In this study, leveraging our group’s established tRNA modification profiling platform (Cao et al., 2020; Yan et al., 2021), we mapped the landscape of tRNA modifications across multiple organs in aging rats. Strikingly, manQ modification exhibited consistent age-dependent depletion in diverse senescence models, emerging as the first tRNA-specific modification demonstrating hallmark-like correlations with aging. This positions manQ decline as a novel biomarker candidate for aging assessment, fulfilling key criteria of biological relevance and dynamic responsiveness. Mechanistic studies revealed that manQ depletion compromises translational fidelity, as demonstrated by aberrant expression of senescence effector proteins such as glycoprotein non-metastatic melanoma protein B (GPNMB). These molecular alterations drive accelerated aging phenotypes through dual mechanisms: error-prone translation and proteostasis network collapse. Crucially, we provide the first experimental evidence that queuine supplementation delays aging and extends healthspan by restoring manQ modification levels, rescuing translational accuracy, and rebalancing proteostatic capacity. Queuine further modulated canonical aging hallmarks, including nutrient-sensing pathways through proteostasis remodeling, enabling systemic regulation of aging. Our findings collectively propose two transformative concepts: (1) tRNA epitranscriptomic signatures constitute a novel class of aging biomarkers, and (2) epitranscriptome-targeted interventions represent a pioneering framework for developing precision geroprotective strategies.

## MATERIALS AND METHODS

### Chemical reagents

Sodium citrate and EDTA anticoagulant blood collection. Sucrose (CAS: 57-50-1, S8271) and agar powder (CAS: 9002-18-0, A8190-500g) were purchased from Beijing Solaibao Biotechnology Co., Ltd. Urea (CAS: 57-13-6, U111902-500g), ammonium persulfate (CAS: 7727-54-0, A112450-500g), TEMED (CAS: 110-18-9, T105496-100mL), NaCl (CAS: 7647-14-5, S433736-500g), EDTA (CAS: 60-00-4, E301914-100mL), HEPES (CAS: 7365-45-9, H109408-500g), KOH (CAS: 1310-58-3, P301749-500g) and paraquat (CAS: 1910-42-5, M106760-100mg) were purchased from Shanghai Aladdin Biochemical Technology Co., Ltd.

Sodium dodecyl sulfate (SDS) (CAS: 151-21-3, S432158-1kg), Tris base (CAS: 77-86-1, T434093-1kg), glycine (CAS: 56-40-6, A111465-2.5kg), NaCl (CAS: 7647-14-5, C111547-500g), KCl (CAS: 7447-40-7, P112143-500g), HCl (CAS: 7647-01-0, H399545-100mL), 2-mercaptoethanol (CAS: 60-24-2, M301574-250mL) and Tween 20 (CAS: 9005-64-5, T434505-100mL) were all purchased from Shanghai Aladdin Biochemical Technology Co., Ltd.

Propionic acid (CAS: 79-09-4, 94425), methyl paraben (CAS: 99-76-3, 1432005), ammonium acetate (CAS: 631-61-8, AX1222), hexafluoroisopropanol (CAS: 920-66-1, 105228-500g), triethylamine (CAS: 121-44-8, 471283-500mL), chloroform (CAS: 67-66-3, 1.02445), formic acid (CAS: 64-18-6, 5.33002, LC-MS grade), methanol (CAS: 67-56-1, 1.06035, LC-MS grade), acetonitrile (CAS: 75-05-8, 1.00029, LC-MS grade), acetic acid (CAS: 64-19-7, 45754), ammonium acetate (CAS: 631-61-8, 5.33004, LC-MS grade) and malic acid (CAS:6915-15-7, M8304) were purchased from Sigma Aldrich (Shanghai) Trading Co., Ltd. MilliQ water was prepared by the ultra-pure water mechanism in the laboratory, with a resistivity of ≥ 18.25MΩ·cm. Queuine hydrochloride (CAS: 69565-92-0, TRC-Q525000-0.5MG), was purchased from Toronto Research Chemicals.

### Biological reagents

Yeast (ATCC9763, LA9380), LB agar powder (L1015-1L) and red blood cell lysate (R1010) were all purchased from Beijing Solaibao Biotechnology Co., Ltd. TRIzol (15596018), acrylamide/Bis 19:1 40% (w/v) solution (AM9022), 2× loading dye (LC6876), SYBR Gold (S11494), RNase T1 (1000U/μL, EN0541), bacterial alkaline phosphatase (2500U, 18011015), DMEM medium (11965092), horse serum (16050122), Opti-MEM medium (31985070), Lipofectamine RNAiMAX transfection reagent (13778150), RIPA lysis buffer (89901), 30% polyacrylamide (R33400) and PVDF transfer membranes (0.45 μm, 88518) were purchased from Shanghai Thermo Fisher Scientific Technology Co., Ltd.

RNase P (10000U, M0660S) was purchased from Beijing Anolun Biotechnology Co., Ltd. SYBR Green qPCR Mix (D7260-25mL), 5× protein loading buffer (P0015L), primary antibody diluent (P0023A-500mL) and horseradish peroxidase-labeled secondary antibody (A0216) were purchased from Shanghai Beyotime Biotechnology Co., Ltd. SA-Biotin (22308-10) was purchased from Suzhou Beaver Biomedical Engineering Co., Ltd. MEM-EBSS (11095080), FBS (10270-106), trypsin (25200-072), penicillin streptomycin glutamine (100×, 10378-016) and PBS (pH=7.4, 10010023) were purchased from Gibco Thermo Fisher Scientific Technology Co., Ltd. cOmplete^TM^ protease inhibitor (EDTA-free, 04693132001) and PhosSTOP calcineurin inhibitors (04906837001) were purchased from Roche Pharmaceuticals in Switzerland. ECL luminescent liquid (180-506) was purchased from Shanghai Tianneng Life Science Co., Ltd. Anti-CDKN2A/p16^INK4a^ antibody (ab270058) was purchased from Shanghai Abcam Trading Co., Ltd. Man2c1 (C-4) antibody (sc-377132, AF790) was purchased from Santa Cruz Biotechnology, Inc. Anti β-actin mouse monoclonal antibody (FI01103S) was purchased from Guangzhou Huijun Biotechnology Co., Ltd.

### Kits

mirVana^TM^ miRNA kit (AM1561) and revertAid^TM^ first strand cDNA synthesis kit (K16225) were purchased from Shanghai Thermo Fisher Scientific Technology Co., Ltd. Senescence associated *β*-Galactosidase staining kit (C0602), BCA protein concentration determination kit (P0011) and glutathione reductase (GSH Px) kit (S0053) were purchased from Shanghai Beyotime Biotechnology Co., Ltd.

The total superoxide dismutase (SOD) kit (A001-3-2) was purchased from Nanjing Jiancheng Bioengineering Institute Co., Ltd. The magnetic bead method blood total RNA extraction kit (DP761) was purchased from Tiangen Biochemical Technology (Beijing) Co., Ltd. The IL-6 ELISA kit (LV30325M) was purchased from Shanghai Aimeng Youning Biotechnology Co., Ltd. Mouse Telomerase (TE) ELISA Kit (ml024416) was purchased from Shanghai Enzyme Linked Biotechnology Co., Ltd.

### Animal origin and ethical approval

SPF-grade male SD rats, aged 2, 6, 24, and 36 months (with 6 rats per age group) and SPF-grade male Balb/c mice, aged 2-6 months were purchased from Zhuhai BaiShiTong Biotech Co., Ltd. SPF-grade male C57BL/6J mice at 16 months (60 mice) were purchased from ZhiShan (Beijing) Health Medical Research Institute Co., Ltd. All animals were housed in the State Key Laboratory of Quality Research in Chinese Medicine at Macau University of Science and Technology (clean-grade animal facility, license number SYXK(Yue)2020-0051). The housing conditions ensured a temperature range of 24-26°C, humidity between 45-65%, and a 12-hour light/dark cycle. Animals had free access to food and water.

All animal experimental protocols were reviewed and approved by the Institutional Animal Care and Use Committee (IACUC) of the State Key Laboratory of Quality Research in Chinese Medicine at Macau University of Science and Technology.

### Cell culture

Human diploid fibroblast 2BS cells, isolated from female fetal lung fibroblast tissue, were purchased from the China Center for Type Culture Collection (CCTCC, NO. GDC0303). The cells were cultured in MEM-EBSS medium supplemented with 10% fetal bovine serum (FBS, Gibco), 100 U/mL penicillin, and 100 μg/mL streptomycin, in a humid atmosphere at 37°C and 5% CO_2_. Based on senescence-associated biomarkers established in our laboratory, cells with a PD of ≤ 30 are classified as young, whereas those with a PD of ≥ 37 are classified as replicatively senescent.

The human embryonic kidney cell line HEK 293T cells were sourced from the Chinese Type Culture Collection Center (CCTCC, NO. GNHu17). Cells were cultured in DMEM medium supplemented with 10% FBS, 100 U/mL penicillin, and 100 μg/mL streptomycin. Other cultivation conditions are the same as those of the 2BS cell line.

### Cultivation of Drosophila melanogaster

The wild-type *D. melanogaster* was provided by China Qidong Fangjing Biotechnology Co., Ltd. They were cultured in standard yeast cornmeal sucrose medium, which consists of 120 mL H_2_O, 12 g cornmeal, 8 g sucrose, 1.5 g dry yeast, 1 g agar powder, 0.5 mL propionic acid, and 0.5 mL 10% nipagin methyl ester). The temperature of the incubator was maintained at (25 ± 1) °C, the humidity at (55 ± 3)%, and the light/dark cycle was 12 h/12 h. Newly emerged *D. melanogaster* were collected within 48 hours under brief CO_2_ anesthesia, fed at a density of 20 per tube, and transferred to fresh culture medium every 3 days throughout the experiment.

### Acute aging model induced by paraquat

Acute aging in mice was induced by intraperitoneal injection of 50 mg/kg paraquat. The survival rate of mice in each group was recorded after modeling. Three days post-injection, animal tissues were collected. The tissues was homogenized and immersed in TRIzol, then stored at −80°C.

### Preparation for queuine detection in plasma

100 µL of acetonitrile was added to 50 µL of plasma to precipitate protein impurities. The mixture was then centrifuged to separate the clarified supernatant, which was subsequently concentrated and dried. The dried supernatant was reconstituted in 20 µL of water.

### Quantitative analysis of queuine using UHPLC-QQQ-MS

The analysis of the queuine base was performed using an Agilent 1290 UHPLC series 6490 triple quadrupole (QQQ) MS system. The Waters ACQUITY UPLC BEH HILIC column (1.7 µm, 2.1 × 100 mm) was used for the separation of queuine. The mobile phase consisted of A (0.2% acetic acid in 10 mM ammonium acetate aqueous solution) and B (acetonitrile with 0.2% acetic acid, 2 mM ammonium acetate, and 0.05 mM malic acid), with a flow rate of 0.4 mL/min. The gradient elution procedure was as follows: 0.00-20.00 min 98-70% B, 20.00-23.00 min 70-60% B, 23.00-26.00 min 60% B, 26.10-30.00 min 98% B. The column temperature was maintained at 40°C, and the injection volume was 1 μL.

The 6490 ESI QQQ-MS was set to positive mode, acquiring data through multiple reaction monitoring (MRM) with *m/z*=278.1→163.1. The instrument parameters applied were as follows: drying gas temperature 200°C, drying gas flow rate 14 L/min, nebulizer pressure 35 psi, sheath gas temperature 350°C, sheath gas flow rate 11 L/min, capillary voltage 3000 V, and nozzle voltage 1500 V. The concentration of queuine in plasma samples was quantified using the external standard method.

### Lifespan

The number of deaths in each group of animals was recorded daily from the experiment’s outset. At the end of the experiment, the time of death for all animals was documented. The survival curve was plotted using GraphPad Prism 7.0, with the horizontal axis representing lifespan (in days or hours) and the vertical axis representing the survival rate.

### Senescence-associated *β*-galactosidase (SA-*β*-gal) staining

According to the kit protocol, a mixture of *β*-galactosidase staining solution and X-Gal substrate was added to the fixed cells or frozen tissue sections of mice and incubated overnight at 37°C. It is worth noting that incubation cannot be carried out in a CO_2_ incubator because carbonate ions can alter the pH of the reaction system.

### RNA extraction and isolation

Total RNA was extracted using TRIzol from cells, human peripheral blood leukocytes, *D. melanogaster*, as well as tissues of rats and mice. According to the experimental protocol, large RNA (>200 nt) and small RNA (<200 nt) were separated from total RNA using the mirVana^TM^ miRNA isolation kit. It should be noted that homogenized tissue needs to be centrifuged in advance to remove debris.

### Real-time fluorescence quantitative PCR (RT-qPCR)

Using the High-Capacity cDNA Reverse Transcription Kit (Roche), 1 μg of large RNA was converted into cDNA. Beyo SYBR^TM^ Green Master Mix (Beyotime) was used for qPCR, which was performed on a Real-Time PCR System (Applied Biosystems). All primer sequence information used in this study is shown in **Table 1**. Relative gene expression was calculated using the 2^-ΔΔCt^ method, normalized against *β*-actin. Each reaction was performed in triplicate, with results presented as mean ± standard deviation (SD).

### Evaluation of healthspan of *D. melanogaster*

#### (a) Lifespan

Newly emerged *D. melanogaster*, collected within 48 hours on standard yeast cornmeal sucrose medium, were randomly divided into a control group and a queuine administration group, each containing 100 flies (50% males and 50% females). The flies were fed at a density of 10 flies per tube, and the culture medium was changed every 3 days. The daily mortality rate of *D. melanogaster* was recorded, and the survival curve was plotted and analyzed using GraphPad 7.0 software.

#### (b) Weight measurement

At each time point (10, 20, and 30 days), ten male and ten female D. *melanogaster* were randomly sampled from each group (control and queuine-treated) and weighed collectively.

#### (c) Exercise

Motor function was assessed using a standard negative geotaxis climbing assay. Briefly, a cohort of flies was transferred into a vertical glass column, and after a 10-minute acclimation period, they were startled by tapping the column to bring them to the bottom. The number of flies that crossed a line marked 8 cm above the bottom within 15 seconds was recorded. The climbing index was calculated as the percentage of flies that successfully crossed the line.

#### (d) Heat stress resistance

Following heat stress treatment (35°C for 30 minutes), flies (50% males and 50% females) were returned to fresh food vials at normal conditions. The number of dead flies was counted after a 24-hour recovery period to determine mortality.

#### (e) Olfactory memory

Two 50 mL centrifuge tubes containing bananas and perforated with small holes were placed in advance inside a 1000 mL beaker. One tube was accessible, allowing D. melanogaster to detect the odor and enter through the holes for feeding, whereas the other tube contained internal obstacles that prevented entry and feeding. For the olfactory memory assay, 20 D. melanogaster (50% males and 50% females), aged 30 days, were used. Prior to testing, flies were placed in empty tubes and starved for 2 hours. They were then transferred to the beaker, and the number of flies that entered the accessible centrifuge tube within 1 minute was recorded.

#### (f) Determination of antioxidant capacity

Equal numbers of male and female D. melanogaster (50% each) were collected and weighed. The flies were thoroughly homogenized in pre-cooled physiological saline. The homogenates were centrifuged at 12,000 r/min at −4 °C to remove tissue debris, and the supernatants were collected for analysis. The activities of superoxide dismutase (SOD) and glutathione peroxidase (GSH-Px) were determined according to the manufacturers’ instructions provided with the respective assay kits

### H&E staining

Mouse tissues were fixed with paraformaldehyde, dehydrated with ethanol, and embedded into sections. The sections were first stained with hematoxylin for 15 minutes. Subsequently, the sections were stained with eosin for 10 seconds. Finally, histological changes were observed under a microscope.

### Extraction of RNA from blood

The Magnetic Bead Total RNA Extraction Kit (DP761) was used for the extraction and purification of RNA from blood. This kit employs specially designed magnetic rods for adsorption and transfer, and finally releases the magnetic beads to obtain high-purity enrichment of RNA.

### Determination of telomerase activity and the content of IL-6

Plasma samples were collected from naturally aging mice. And the telomerase detection kit and IL-6 enzyme linked immunosorbent assay (ELISA) kit were used to quantify telomerase activity and the content of IL-6 in plasma, according to standard operating procedures.

### Open field test

Mice were individually placed in open-field boxes (40 cm × 40 cm × 40 cm) and allowed to explore freely for 7 minutes. Locomotor activity during the last 5 minutes of the test was recorded and analyzed using Tracker software. The following parameters were measured: total movement trajectory, open-field score (number of grid crossings), time spent in the central area (20 cm × 20 cm), and time spent in the peripheral area. After each trial, the open-field apparatus was cleaned with 75% ethanol to eliminate residual odors and prevent interference with subsequent tests.

### Object Location Test (OLT) and Novel Object Recognition Test (NORT)

Objects A and B were placed on the same side of the open-field box (40 cm × 40 cm × 40 cm). The exploration zone was defined as a circular area with a radius of 2 cm centered on each object. The time spent exploring objects A and B was recorded. For the OLT, object B was relocated to the opposite side of the open-field box, and the change in the proportion of exploration time for objects in different spatial positions was calculated. For the NORT, object B was replaced with a novel object (object C), and the percentage of time spent exploring the novel object was determined.

### Rotarod test for motor coordination and fatigue

Three days prior to formal testing, mice in the control (2M), natural aging (blank, 18M), and queuine-treated groups (18M) were trained daily on a rotating rod apparatus. During the formal test, the latency to fall (residence time) at 15 r/min was recorded.

### Grip strength test

Muscle strength was assessed using a grip force meter. Each mouse was tested three times, and the highest value was recorded as the maximal grip strength.

### Y-Maze Test

Mice from the control (10M), natural aging (blank, 26M), and queuine-treated groups (26M) were placed individually in a Y-maze consisting of three identical arms (30 cm × 6 cm × 15 cm) and allowed to explore freely for 5 minutes. Movement trajectories and spontaneous alternation behavior were recorded using the TopScan 3.0 system.

### DNA extraction from mouse blood, feces, and tissues

DNA was extracted and purified from mouse blood, fecal microbiota, and tissues using a DNA extraction kit (DP304) according to the manufacturer’s instructions. The kit utilizes magnetic bead technology for nucleic acid adsorption and transfer, enabling high-purity DNA enrichment.

### Agarose gel electrophoresis

DNA or RNA concentration and purity were assessed using a NanoDrop spectrophotometer (Thermo Fisher Scientific). A total of 500 ng of DNA or RNA was mixed with an equal volume of 2× loading buffer (95% formamide, 18 mM EDTA, 0.025% SDS, xylene blue, and bromophenol blue) and analyzed by 1% agarose gel electrophoresis.

### DNA methylation clock analysis

DNA was extracted from mouse white blood cells and subjected to bisulfite sequencing. Sequencing reads were processed using Illumina base-calling software and aligned to the reference genome using BSMAP. DNA methylation levels at CpG sites were quantified as the ratio of reads supporting cytosine (C) to the total reads supporting C and thymine (T). DNA methylation age was estimated using a previously established elastic net regression model and analytical pipeline described by Cao et al.

### Hematological and Blood Biochemical Analysis

Peripheral blood (20 μL) was collected into anticoagulant tubes. Complete blood counts were performed using an automated veterinary hematology analyzer (BC-2800vet, Mindray). Plasma was separated for biochemical analyses, including liver function, renal function, myocardial enzymes, blood glucose, and lipid profile, using an automated biochemical analyzer (Chemray 800).

### 16S rRNA sequencing

Total DNA was extracted from fecal samples of control mice (6M), aging mice (blank, 22M), and queuine-treated mice (22M). The 16S rRNA gene was amplified using the primers: forward primer 5’-ACTCCTACGGGAGGCAGCA-3’ and the reverse primer 5’-GGACTACHVGGGTWTCTAAT-3’. Amplicons were sequenced using paired-end sequencing on the Illumina platform.

### GC-MS/MS analysis of short-chain fatty acids (SCFAs) in mouse fecal

50 mg of mouse fecal sample was mixed with 0.2 mL of phosphoric acid (0.5% v/v) solution and vortexed for 10 minutes. 500 μL of methylation tert-butyl ether (MTBE) solvent containing an internal standard was added. Samples was sonicated in an ice bath for 5 minutes and centrifuged at 12000 r/min at 4°C for 10 minutes. The upper layer (200 μL) was collected for GC-MS/MS analysis.

An Agilent 7890B-7000D GC-MS/MS instrument equipped with a DB-FFAP column (30 m × 0.25 mm × 0.25 μm) was used to quantify the levels of six SCFAs in the samples. The instrument parameters were set as follows: chromatographic separation was in split mode 1:1, carrier gas was helium, flow rate was 1.2 mL/min, injection volume was 2 μL, column box temperature program was 90°C hold on for 1 minute, raised to 100°C at a rate of 25°C/min, raised to 150°C at a rate of 20°C/min, hold on for 0.6 min, raise to 200°C at a rate of 25°C/min, hold on for 0.5 min, after running for 3 min. The inlet temperature was 200°C, the transfer line temperature was 230°C, the ion source temperature was 230°C, and the fourth stage rod temperature was 150°C.

### Urea-PAGE electrophoresis

RNA concentration and purity were measured using a NanoDrop spectrophotometer. A total of 200 ng RNA or 50 ng purified tRNA was mixed with 2× loading buffer and analyzed by 6% urea-PAGE.

### RNaseT1 digestion

Total tRNA was denatured at 95°C for 3 minutes and immediately chilled on ice. Ten micrograms of RNA were digested with RNase T1 (50 U/μg) in 220 mM ammonium acetate at 37°C for 2.5 hours. The reaction was terminated at 70°C for 10 minutes and diluted to 50 μL with ion-pair mobile phase A for tRNA mapping.

### UHPLC-QTOF-MS analysis for tRNA mapping

UHPLC-QTOF-MS analysis of tRNA fragments was performed as previously described (Pan et al., 2021). Briefly, enzymatically digested tRNA fragments were analyzed using an Agilent 1290 UHPLC system coupled with an Agilent 6545 Q-TOF MS spectrometer. Chromatographic separation was achieved on an ACQUITY UPLC OST C18 column using an ion-pair mobile phase system optimized for oligonucleotide analysis. Mass spectrometric detection was performed in negative ion mode with auto MS/MS acquisition for structural characterization. The mass accuracy between theoretical and experimentally observed masses was controlled within 5 ppm to ensure reliable fragment identification. All experiments were conducted in triplicate, and data are presented as mean ± standard deviation (SD)

### Purification of individual tRNAs

Specific tRNAs, including tRNA^Asp(manQUC)^, tRNA^Asn(QUU)^, tRNA^His(QUG)^, and tRNA^Tyr(galQΨA)^, were purified from small RNA fractions using a biotinylated DNA-probe based affinity capture method, as previously described (Ren et al., 2023). To enhance purification efficiency and specificity, two complementary biotinylated DNA probes were designed for each target tRNA, corresponding to sequences at the 5’ and 3’ termini (**Table 2**). Briefly, small RNA samples were incubated with the corresponding biotinylated DNA probes at 65°C for 2 h to facilitate hybridization. The resulting biotinylated DNA–tRNA hybrids were subsequently captured using Pierce™ streptavidin-coated magnetic beads through high-affinity biotin–streptavidin interactions. After extensive washing to remove non-specifically bound RNA, the target tRNAs were released by heat denaturation in RNase-free water at 70°C and collected for downstream analyses.

### Nucleoside preparation and manQ quantification

To prepare total nucleosides for tRNA modification analysis, 3 μg of tRNA was incubated in a reaction buffer containing 20 mM MgCl_2_, 80 mM Tris-HCl (pH 8.0), 3U phosphodiesterase I, 3U bacterial alkaline phosphatase. The total reaction volume was adjusted to 20 μL with RNase-free water and incubated at 37°C for 8 h to achieve complete enzymatic digestion.

Nucleoside quantification was performed using an Agilent 1290 UHPLC coupled with 6490 triple quadrupole (QQQ) mass spectrometer. Chromatographic separation of tRNA modifications was achieved using an Agilent Poroshell 120 EC-C18 column (2.7 µm, 4.6×100 mm). The mobile phase consisted of 0.1% formic acid (A) and acetonitrile (B), with a flow rate of 0.4 mL/min. The gradient program was used as follows: 0.00-2.50 min 1.5% B, 2.50-6.00 min 1.5%-4%, 6.00-12.00 min 4%-15%, 12.00-16.00 min 15%-45%, 16.00-20.00 min 45%, 20.00-23.00 min 80%, 23.00-24.00 min 1.5%, 24.00-25.00 min 1.5%. The column temperature was set at 30°C, and the injection volume was 2 μL.

The 6490 ESI-QQQ-MS was operated in positive ion mode. Multiple reaction monitoring (MRM) transitions were set as follows: manQ, *m/z* 572.0→163.0; guanosine, *m/z* 284.0→152.0. Other instrument parameters were identical to those described for the quantitative analysis of queuine. The peak area of guanosine was used as an internal reference for data normalization.

### mRNA transcriptome analysis

Total RNA was extracted and enriched for mRNA using NEB Next® Poly(A) mRNA Magnetic Isolation Module (NEB, USA). Libraries were prepared using the NEBNext Ultra RNA Library Prep Kit and sequenced on the Illumina platform (paired-end 150 bp). Clean reads were aligned to the mouse genome (mm10) using HISAT2. Differential expression analysis was performed using DESeq2 with Benjamini-Hochberg correction. Genes with fold change >2 and adjusted p < 0.05 were considered significant. All the differentially expressed genes were used for heat map analysis and KEGG ontology enrichment analyses. For KEGG enrichment analysis, a P-value < 0.05 was used as the threshold to determine significant enrichment of the gene sets.

### Molecular docking

The 2D structure of queuine was retrieved from PubChem and optimized using Chem3D. The JAK2 crystal structure (PDB ID: 7F7W) was obtained from the RCSB PDB database. Docking simulations were performed using AutoDock Tools with semi-flexible docking. The lowest binding energy conformation was visualized using PyMOL.

### Gene silencing and overexpression

siRNA targeting *MAN2C1* was purchased from Suzhou Beixin Biotechnology Co., Ltd. The siRNA sequences were as follows: sense strand (5’-3’), CAUGAUGUGGUGACUGGAAGCUGCA; the antisense strand (5’-3’), UGCAGCUUCCAGUCACCACAUCAUG. Transient transfection was performed using Lipofectamine™ RNAiMAX (Invitrogen) in accordance with the manufacturer’s instructions. Cells were transfected with the indicated siRNA under optimized conditions and subsequently harvested at the specified time points for downstream analyses.

For overexpression, the *MAN2C1* gene was cloned into the pEZ-Lv105 expression vector and packaged into lentiviral particles using a lentiviral production system (System Biosciences, USA) according to the manufacturer’s instructions. 2BS cells were transduced with the recombinant lentivirus, and stable cell lines were established by selection with puromycin. Overexpression efficiency was validated by qPCR. The primer sequences were as follows: forward, 5’-GCGGTAGGCGTGTACGGT-3’; reverse, 5’-ATTGTGGATGAATACTGCC-3’.

### Western blot (WB)

Cells or tissues were lysed in RIPA buffer supplemented with protease and phosphatase inhibitors. Protein lysates were denatured at 95°C for 10 min, and equal amounts of protein (30 μg per sample) were separated by 10% SDS–PAGE and subsequently transferred onto polyvinylidene fluoride (PVDF) membranes. Membranes were blocked with 5% skim milk and incubated overnight at 4°C with primary antibodies against Man2c1 (1:1000 dilution, Santa Cruz Biotechnology), p16 (1:1000 dilution, Abcam), or β-actin (1:5000 dilution, Guangzhou Huijun Biotechnology Co., Ltd.). After washing, membranes were incubated with horseradish peroxidase (HRP)-conjugated secondary antibodies. Protein bands were visualized using an enhanced chemiluminescence detection system (Tanon).

### Gene knockout

The *GTDC1* gene in 2BS cells was knocked out using CRISPR/Cas9-mediated gene editing delivered via electroporation. The guide RNAs (gRNAs) used as follows: gRNA-A1, 5’-CTCTCCTGACATTTCTTGAC-AGG-3’; and gRNA-B1: 5’-AGATCCTTGGGTCTGTGATC-AGG-3’. After electroporation, the *GTDC1* gene knockout cell pool was established and validated through PCR and sequencing. The primer used for PCR identification and sequencing are listed in **Table 3**.

### Differential proteomic analysis

Proteins were enzymatically digested with trypsin to generate peptides. The resulting peptide mixtures were desalted using C18 solid-phase extraction cartridges, lyophilized, and reconstituted in 40 μL of 0.1% formic acid. Peptide concentrations were determined by measuring ultraviolet absorbance at 280 nm (OD280) with a NanoDrop spectrophotometer. Equal amounts of peptides were subjected to data-independent acquisition (DIA)–based proteomic analysis using an Astral high-resolution mass spectrometer coupled to a Vanquish Neo nano-flow UHPLC system (Thermo Scientific) for chromatographic separation. The mass spectrometer was operated in positive ion mode. MS1 spectra were acquired over an m/z range of 380–980 at a resolution of 240,000 (at m/z 200), with a normalized automatic gain control (AGC) target of 500% and a maximum injection time (IT) of 5 ms. MS2 spectra were acquired in DIA mode using 299 isolation windows with a width of 2 m/z. Peptide fragmentation was performed by higher-energy collisional dissociation (HCD) at a collision energy of 25 eV. For MS2 acquisition, the normalized AGC target was set to 500%, and the maximum IT was 3 ms. Raw DIA data were processed using DIA-NN software. Database searches were conducted with trypsin specified as the proteolytic enzyme, allowing up to one missed cleavage. Carbamidomethylation of cysteine was defined as a fixed modification, whereas oxidation of methionine and protein N-terminal acetylation were set as variable modifications. Protein identifications were filtered at a false discovery rate (FDR) of <1%.

Functional annotation of the identified proteins was performed using Blast2GO (BLASTP 2.8.0+), including sequence alignment, Gene Ontology (GO) term mapping, GO annotation, and InterProScan-based annotation enhancement. Kyoto Encyclopedia of Genes and Genomes (KEGG) pathway annotation was carried out using KOBAS 3.0. Enrichment analyses of GO terms and KEGG pathways were performed using Fisher’s exact test, comparing the annotation distribution of the target protein set with that of all identified proteins as the background. Kinases were further annotated and analyzed using publicly available kinase-related databases, including PhosphoSitePlus (PSP) and Phospho.ELM.

### Targeted metabolomics analysis

To extract metabolites, 400 μL of cold methanol/acetonitrile (1:1, v/v) was added to plasma samples to precipitate proteins and extract metabolites. Stable-isotope internal standards were added to the extraction solvent for absolute quantification. The mixture was vortexed thoroughly and centrifuged at 14,000 g for 20 min at 4°C. The supernatant was collected and dried in a vacuum centrifuge. The dried extracts were reconstituted in 100 μL of acetonitrile/water (1:1, v/v), centrifuged again at 14,000 g for 15 min at 4°C, and the final supernatant was used for LC–MS analysis.

Metabolite analysis was performed using an Agilent 1290 Infinity UHPLC system coupled to a 6500+ QTRAP mass spectrometer (AB Sciex). Separation was carried out on both a HILIC column (Waters UPLC BEH Amide, 2.1 × 100 mm, 1.7 μm) and a C18 column (Waters UPLC BEH C18, 2.1 × 100 mm, 1.7 μm). For HILIC separation, the column temperature was set at 35°C and the injection volume was 2 μL. Mobile phase A consisted of 90% water with 2 mM ammonium formate and 10% acetonitrile, and mobile phase B was 0.4% formic acid in acetonitrile. The gradient was as follows: 85% B (0-1 min), 80% B (3-4 min), 70% B (6 min), 50% B (10–15.5 min), and 85% B (15.6–23 min), at a flow rate of 300 μL/min. For reversed-phase (RPLC) separation, the column temperature was 40°C and the injection volume was 2 μL. Mobile phase A was 5 mM ammonium acetate in water, and mobile phase B was 99.5% acetonitrile. The gradient was 5% B (0 min), 60% B (5 min), 100% B (11–13 min), and 5% B (13.1–16 min), at a flow rate of 400 μL/min. Samples were maintained at 4°C throughout the analysis.

Quantitative data were acquired using multiple reaction monitoring (MRM), and the MRM transitions are listed in **Table S1**. Quality control (QC) samples were included in the run sequence to monitor system stability and repeatability. The 6500+QTRAP was operated in positive/negative ion switching mode. For positive electrospray ionization (ESI), the source temperature was 580°C, GS1 was 45, GS2 was 60, curtain gas was 35, and the ion spray voltage was +4500 V. For negative ESI, the same settings were used except that the ion spray voltage was −4500 V.

### Data analysis

All experiments were performed with three biological replicates, and data are presented as the mean ± standard deviation (SD). Statistical analyses were conducted using one-way analysis of variance (ANOVA) for multiple group comparisons. Statistical significance was defined as p < 0.05 (*), p < 0.01 (**), and p < 0.001 (***). Sequence denoising was carried out using QIIME2 (version 2019.4), Vsearch (v2.13.4_linux_x86_64), and Cutadapt (v2.3). Alpha diversity, beta diversity, microbial community composition, differential species analysis, and biomarker identification were comprehensively analyzed using RStudio software. GraphPad Prism 7.0, SIMCA 14, ChemDraw 20, and Figdraw 2.0 were used for statistical visualization, chemical structure illustration, and schematic diagram preparation.

## RESULTS

### Age-associated decline of circulating queuine levels in mammals

To quantify queuine dynamics in plasma during aging, we performed ultra-high-performance liquid chromatography coupled with triple quadrupole mass spectrometry (UHPLC-QQQ-MS) using a validated external standard method. The diagnostic transition *m/z* 278.0→163.0, corresponding to queuine’s characteristic fragmentation pattern (**Fig. S2A**), was selected for quantification. Longitudinal analysis revealed progressive depletion of plasma queuine in rats from 18.22 ± 2.72 ng/mL (6 months) to 8.83 ± 0.75 ng/mL (36 months), showing strong age correlation (slope = −0.3225, *p*<0.0001; **Fig. 1A**). Human cohort analysis mirrored this trend, with plasma levels declining by over 65% between young adults (30-40 years) and older individuals (> 60 years) (slope = −0.05188, *p*<0.0001; **Fig. 1B**). Notably, this depletion may be exacerbated by age-related inflammation. Elevated interferon production in older adults, a hallmark of inflammaging, could impair queuine utilization, as demonstrated by Elliott et al. who identified interferon-mediated inhibition of queuine uptake mechanisms (Elliott and Crane, 1990). This dual regulation likely synergistically drives tRNA modification deficits during aging.

**Figure 1.**
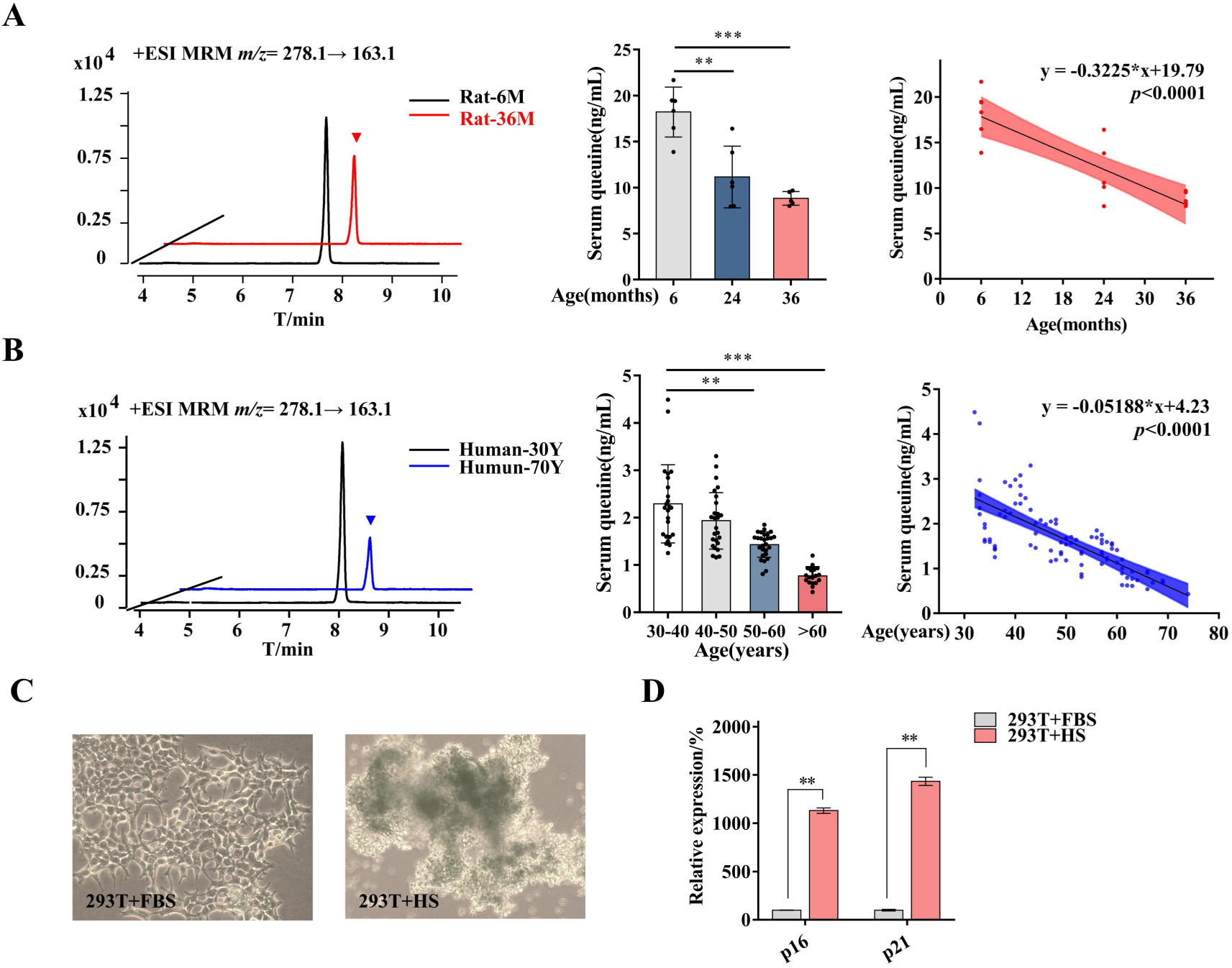
Age-associated queuine depletion correlates with cellular senescence. (A) Age-dependent decline of serum queuine in rats. Queuine concentrations in male rats at 6, 24, and 36 months (n = 6/group; \*\*\**p* < 0.0001 by linear regression). (B) Age-dependent decline of plasma queuine in humans. Queuine levels across age cohorts: 30-40, 40-50, 50-60, and ≥60 years (n = 25/group; \*\*\**p* < 0.0001 by linear regression). (C) Senescence-associated *β*-galactosidase (SA-*β*-gal) activity. Representative images of SA-*β*-gal staining (blue) in HEK 293T cells cultured in horse serum (HS) vs. fetal bovine serum (FBS) for 72 hr (scale bar: 50 μm). (D) Senescence marker expression. Relative mRNA levels of p16 and p21 in HS-vs. FBS-cultured cells (n = 3; \**p* < 0.05, \*\**p* < 0.01, \*\*\**p* < 0.001).

Comparative profiling revealed significantly lower queuine levels in horse serum (HS) versus fetal bovine serum (FBS) (**Fig. S2B**). HEK293T cells maintained in HS exhibited pronounced senescence phenotypes, including morphological deterioration, senescence activation and cell cycle arrest markers. Reduced cell adhesion capacity observed via phase-contrast microscopy. Senescence-associated *β*-galactosidase positive cells was significantly increased (*p*<0.001 vs. FBS controls, **Fig. 1C**). Upregulation of p16^INK4a^ (cyclin dependent kinase inhibitor 2A, CDKN2A) and p21^WAF1/CIP1^ (cyclin dependent kinase inhibitor 1A, CDKN1A) mRNA levels (10.8-and 14.2-fold increases, respectively; *p*<0.01, **Fig. 1D**) were also observed. This serum-dependent senescence phenotype strongly correlates with queuine availability, suggesting its depletion may drive cellular aging.

### Queuine supplementation extends lifespan across multiple dimensions

To systematically evaluate the effects of queuine on lifespan across multiple species and dimensions, we examined naturally senescent cells, *Drosophila melanogaster*, paraquat-induced acute aging models, and naturally aging mice. Initially, to address queuine deficiency in horse serum (HS), we supplemented it with 10 ng/mL queuine (HS+Q). Young 2BS cells (30th generations) were cultured in three media: FBS, HS, and HS+Q (**Fig. 2A**). Similar to observations in HEK 293T cells, HS-cultured 2BS cells displayed reduced adhesion and elevated senescence markers, particularly p21 (**Fig. 2B**). Crucially, queuine supplementation fully rescued this senescence phenotype, confirming queuine’s role in counteracting serum deficiency-induced aging. We next extended this approach to FBS, supplementing it with 10 ng/mL queuine (2× endogenous FBS levels; 6.92 ng/mL baseline) across serial passaging (generations 30-39th) (**Fig. 2C**). Aged 2BS cells (39th generations) in queuine-enriched FBS exhibited marked suppression of senescence markers compared to controls (p16: −74.64%, p21: −71.21%; **Fig. 2D**). Dose-response analysis revealed concentration-dependent effects: high-dose supplementation (50 ng/mL) further reduced p16 and p21 expression by 253.89% and 77.63%, respectively, in late-passage cells (**Fig. 2E-F**). This demonstrates a significant enhancement of anti-aging efficacy with elevated queuine concentrations (*p*<0.01 vs. 10 ng/mL group).

**Figure 2.**
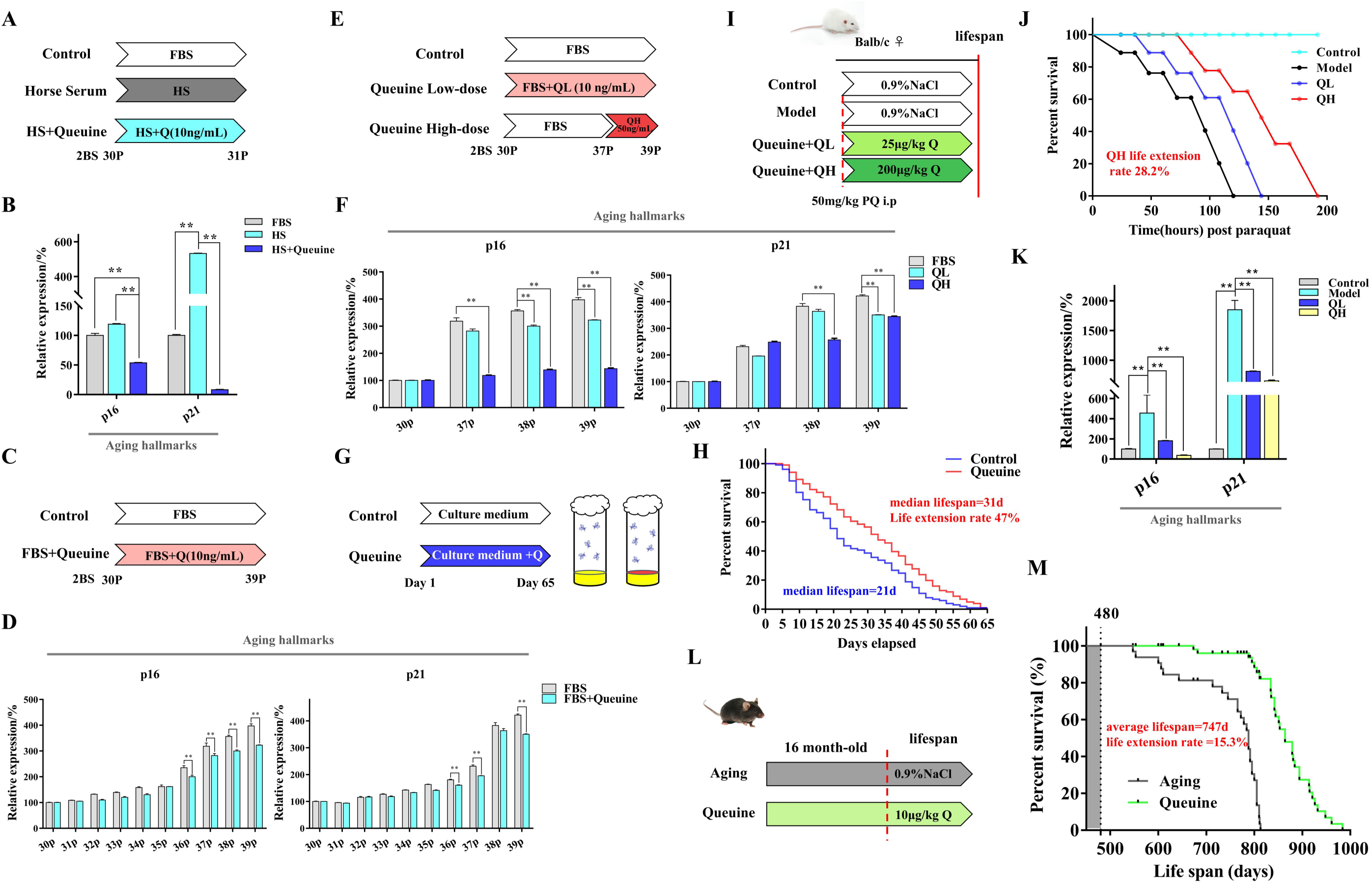
Queuine supplementation attenuates senescence and extends survival across models. (A) Experimental design: Queuine rescue in HS-cultured 2BS cells. Schematic of early-passage (P30) cells treated with: FBS (control), HS (queuine-deficient), or HS + 10 ng/mL queuine (HS+Q). (B) Senescence marker reversal. p16 and p21 mRNA in HS-cultured 2BS cells ± queuine rescue (n=3; \*\**p*<0.01 vs. HS). (C) Experimental design: Queuine in natural aging. 2BS cells (P30) cultured continuously in FBS ± 10 ng/mL queuine. (D) Natural senescence attenuation. Senescence markers in P39 2BS ± queuine (n=3; \*\**p*<0.01 vs. FBS). (E) Dose-response design. 2BS cells treated with 0/10/50 ng/mL queuine in FBS. (F) Dose-dependent effects. Marker expression at different doses (n=3; \*\**p*<0.01 vs. 0 ng/mL). (G) *D. melanogaster* supplementation protocol. Queuine (50 ng/mL) added to food. (H) *D. melanogaster* lifespan extension. Survival curve of queuine treatment and control flies (n=200; log-rank *p*<0.001). (I) Acute mouse aging protocol. Paraquat (50 mg/kg) ± queuine (25/200 μg/kg) in 6-month C57BL/6J males. (J) Acute survival rescue. Percent survival post-paraquat ± queuine (n=6/group; \*\**p*<0.01). (K) Senescence markers. p16 and p21 mRNA in acute model (n=3; \**p*<0.05 vs. paraquat-only). (L) Chronic mouse aging protocol. Queuine (10 μg/kg) administered orally to 16-month C57BL/6J males. (M) Longevity extension. Kaplan-Meier curve of naturally aged mice (n=30; log-rank *p*<0.001).

The evolutionary conservation of queuine metabolism, encompassing uptake, integration, and utilization pathways, across diverse eukaryotes prompted us to investigate its anti-aging effects in *D. melanogaster*. Using a bottle-feeding model (**Fig. 2G**), we observed that queuine supplementation (50 ng/bottle) significantly extended both median and maximum lifespans compared to controls. While control flies exhibited a median lifespan of 21 days and maximum longevity of 59 days, queuine-treated cohorts showed 47.46% extension in median survival (31 days) and 10.17% increase in maximum lifespan (65 days) (**Fig. 2H**). Strikingly, these improvements occurred without observable toxicity, as evidenced by maintained feeding behavior and mobility in treated groups.

The paraquat-induced acute aging model provides accelerated assessment of anti-aging interventions through oxidative stress mechanisms mimicking natural aging(Singh et al., 2023). We administered oral queuine at two doses (QL: 25 μg/kg; QH: 200 μg/kg) to Balb/c mice following paraquat challenge (**Fig. 2I**). Dose-dependent survival improvements emerged: QH extended mean survival to 100 hours (QL 84 h, QH 100 h vs. model group 78 h; *p*<0.05) (**Fig. 2J**). Tissue analyses revealed profound suppression of aging markers in QH, p16 was reduced by 2.7-fold, accompanied by p21 reductions of 2.3-fold compared to the model controls (**Fig. 2K**). In age-matched C57BL/6 males (16-month-old, human-equivalent 50 years) receiving 10 μg/kg queuine every three days (**Fig. 2L**), treatment conferred remarkable survival advantage: Queuine extended mean lifespan from 747.13 days (controls) to 861.13 days (15.3% increase) (*p*<0.001, **Fig. 2M**). The median survival increased from 788 days in controls to 864 days in the treated group, representing a 9.64% extension in lifespan. Additionally, maximal survival was extended by 141 days (from 813 to 984 days), and the hazard ratio was 0.19 (95% CI: 0.09-0.38). This conserved longevity enhancement, spanning cellular models (2BS), invertebrates (*D. melanogaster*), and mammals, establishes queuine as a multimodal anti-aging agent.

### Queuine supplementation enhances healthspan across biological scales

While lifespan extension provides fundamental validation, we further established queuine’s healthspan benefits through multidimensional assessment in three model systems. Our integrated analysis combined behavioral phenotyping, histopathology, biochemical profiling, and multi-parametric functional testing across *D. melanogaster*, acute aging mice, and natural aging mice (**Fig. 3A**). In *D. melanogaster*, longitudinal monitoring revealed queuine’s capacity to decouple chronological aging from functional decline. Although treatment showed no impact on body mass (**Fig. 3B**), it conferred striking protection against age-related functional deterioration. By day 30 post-eclosion (equivalent to human late adulthood), queuine-treated flies exhibited: 11.2% greater climbing capacity in negative geotaxis assays (*p*<0.05; **Fig. 3D**) 2.3-fold extended survival under thermal stress (42°C) (*p*<0.001; **Fig. 3E**), 48.5% improvement in olfactory memory retention *p*<0.001; **Fig. 3F**), and 53.9% reduction in reactive oxygen species (vs. age-matched controls; *p*<0.001; **Fig. 3G**).

**Figure 3.**
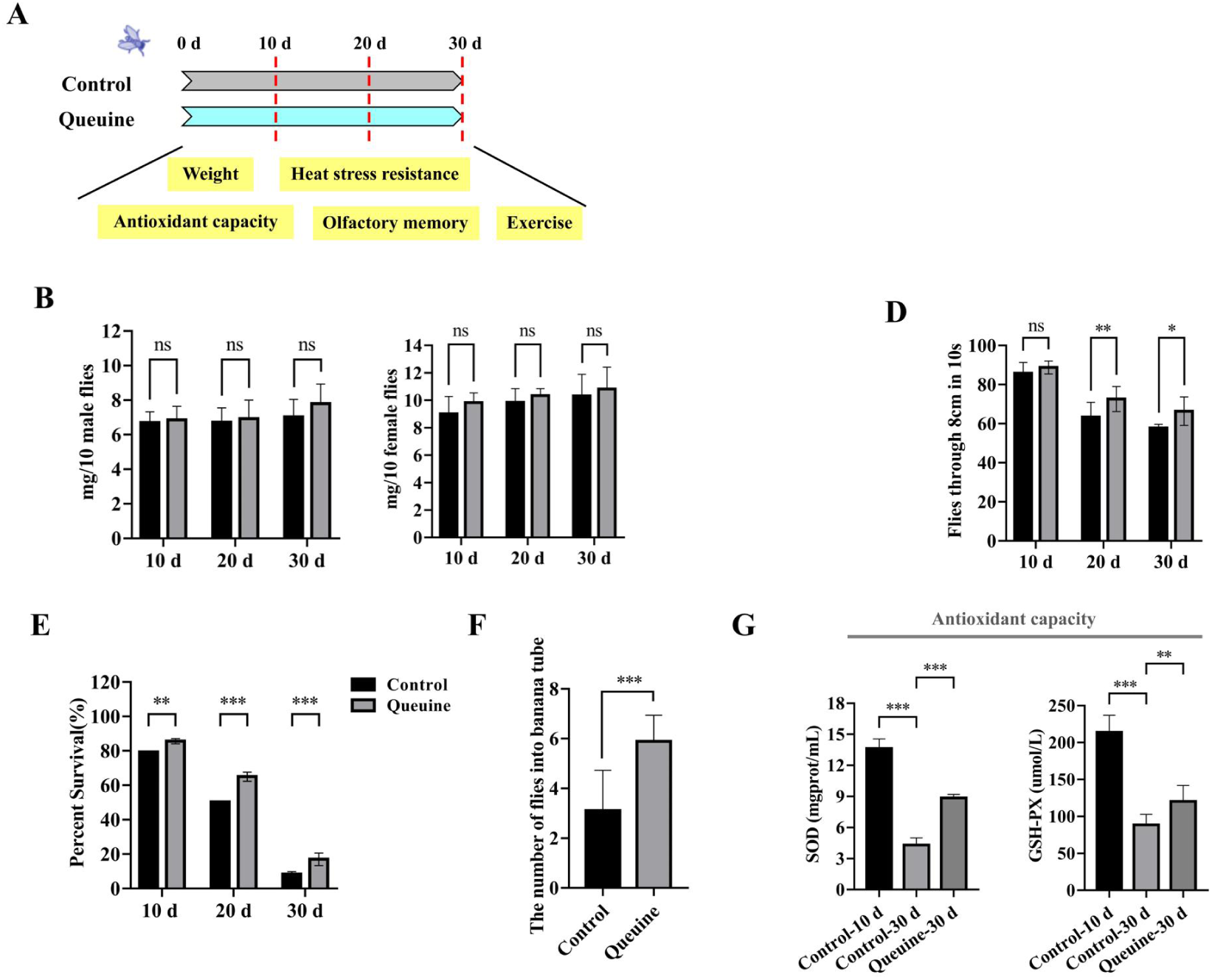
Queuine supplementation extends healthspan in *D. melanogaster*. (A) Schematic of experimental design assessing queuine-mediated healthspan improvement in *D. melanogaster*. Wild-type flies and flies supplemented with queuine (50 ng per food bottle) were evaluated for weight, exercise capacity, heat stress resistance, olfactory memory, and antioxidant capacity at 10, 20, and 30 days post-eclosion. (B-G) Longitudinal analysis of healthspan parameters in control and queuine-supplemented flies (n=10 cohorts): body weight (B), exercise performance (D), heat stress survival (E), olfactory memory retention (F), and systemic antioxidant activity (G).

We conducted longitudinal monitoring of 16-month-old C57BL/6 males (human-equivalent 50 years) receiving 10 μg/kg queuine once every three days, with age-matched controls and young controls. Multi-parametric assessments were performed at 8, 24, and 40 weeks post-treatment (**Fig. 4A**). Following a period of two months during which the mice were administered the experimental treatment, a significant reduction in the levels of the aging hallmarks p16 and p21 (**Fig. 4B**) and the inflammatory factor IL-6 (**Fig. 4D**) was observed in the leukocytes of the treated mice when compared with the levels in the control group (2-month-old). Concurrently, telomerase activity (**Fig. 4C**) and antioxidant capacity (**Fig. 4E**) exhibited a marked enhancement in the treated mice. Furthermore, the behavioral experiments revealed that mice treated with queuine exhibited a substantial enhancement in open field scores, demonstrating superior performance in terms of free exploration (**Fig. 4F**), anti-fatigue (**Fig. 4I**), and recognition ability for novel locations and objects (**Fig. 4H**), in comparison to the blank group of mice.

**Figure 4.**
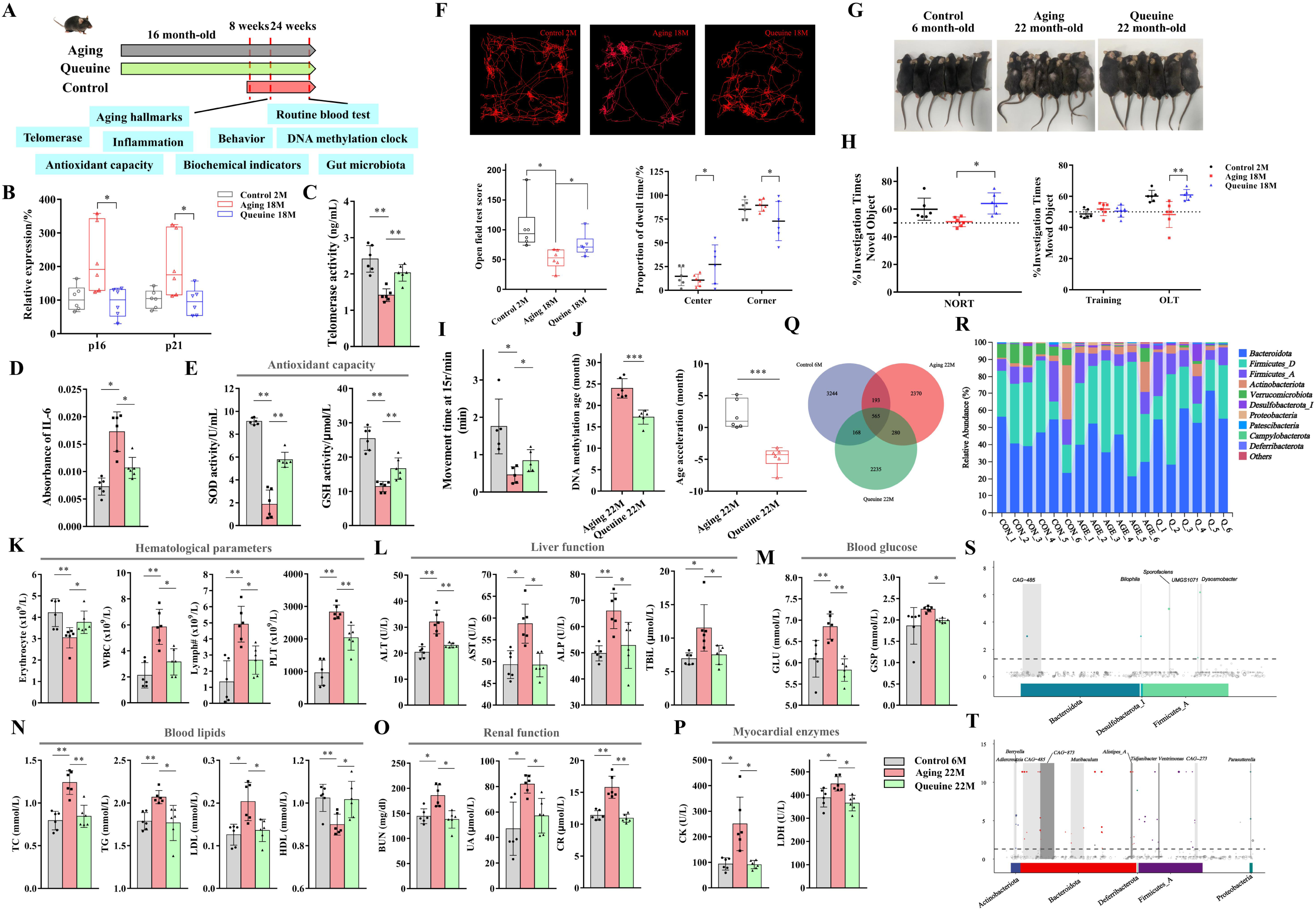
Multisystem rejuvenation through queuine supplementation in naturally aging mice. (A) Experimental design: Middle-aged (16-month-old) wild-type male C57BL/6J mice received oral queuine (10 μg/kg bw/day; n=6) or vehicle once every three days. Aging hallmarks, behavioral/cognitive function, molecular biomarkers (telomerase activity, DNA methylation clock, inflammatory cytokines, antioxidant capacity), clinical biochemistry (hematology, liver/kidney function, glucose/lipids, myocardial enzymes), and gut microbiota were assessed at 8, 24, and 40 weeks post-treatment. (B-P) Queuine-mediated improvements in: Molecular aging biomarkers: p16/p21 expression (B), telomerase activity (C), serum IL-6 (D), systemic antioxidant capacity (E). Behavior/cognition: Open Field locomotion (F), physical appearance (G), Novel Object Recognition (H-left) and Object Location Memory (H-right), endurance (I). Epigenetic aging: DNA methylation age (J). Clinical pathology: Hematological indices (K), liver transaminases (L), fasting glucose (M), lipid profile (N), renal biomarkers (O), and cardiac enzymes (P). (Q-T) Gut microbiota modulation by queuine: (Q) Venn diagram of core microbial OTUs in young (6mo), aged (22mo), and aged+queuine (22mo) mice. (R) Phylum-level taxonomic composition, (S) Species significantly enriched in aged+queuine vs. aged mice. (T) Species significantly enriched in aged vs. young mice.

Following a six-month queuine treatment regimen, significant improvements in hair color were observed in treated mice (**Fig. 4G**). Concurrently, substantial increases in red blood cells and notable decreases in white blood cells and lymphocytes were detected (**Fig. 4K**). Biochemical indicators, encompassing liver function (**Fig. 4L**), renal function (**Fig. 4O**), blood glucose (**Fig. 4M**), blood lipids (**Fig. 4N**), and myocardial enzyme profiles (**Fig. 4P**), also showed significant improvement. Most strikingly, queuine treatment over six months reduced the DNA methylation age of the treated group by 4.72 months compared to controls (**Fig. 4J**).

Analysis of gut microbiota via 16S rRNA sequencing in young control, aging blank control, and queuine-treated mice revealed alterations in α-diversity, β-diversity, species composition, and species differences (**Fig. S4A**). The treatment group exhibited an increased α-diversity index (Pielou_e, *p*=0.036), greater community homogeneity, and broader rank-abundance curves (indicating spectral broadening) (**Fig. S4B**). β-diversity analyses (PCoA and NMDS) demonstrated enhanced microbial stability in queuine-treated mice (**Fig. S4C**). Significant differences in species composition were evident among the three groups (**Fig. 4Q**). Specifically, the abundance of *Akkermansia* (a beneficial genus within the Verrucomicrobiota phylum) decreased in aged mice, while Proteobacteria phylum abundance was upregulated in the aging blank control group. Conversely, the abundance of *Lactobacillus* (a beneficial genus within the Firmicutes_D phylum) was significantly upregulated in queuine-treated mice (**Fig. 4R, S4D**). Species difference analysis indicated that the microbiota of queuine-treated mice was more stable and compositionally resembled that of young mice (**Fig. S4E**), with signature species phylogenetically closer to those in young mice (**Fig. S4F**). In contrast, potentially harmful genera (*Allobaculum*, *Eubacterium*, *Parasutterella*) constituted predominant bacterial groups in the aging blank control group, showing increased abundance (**Fig. 4T, S4G**). Queuine treatment, however, led to a significant increase in beneficial bacteria, including *Lactobacillus*, *Rikenellaceae* (family), *Limosilactobacillus*, *Sporofaciens*, and *Deferribacterales* (order) within the intestinal microbiota (**Fig. 4S, S4G**) (O’Toole and Jeffery, 2015). Furthermore, fecal concentrations of all six measured short-chain fatty acids (SCFAs) were significantly elevated in the treatment group compared to the blank control group (**Fig. S4H**), indicating a potential enhancement in intestinal health for aged mice.

After 10 months of queuine treatment, mice exhibited significantly enhanced muscle strength compared to untreated controls (**Fig. S5B**). The Y-maze test revealed augmented spontaneous exploratory behavior in treated mice (**Fig. S5A**). Furthermore, *β*-galactosidase staining of multiple organs showed significantly fewer senescence-associated positive (blue) cells in queuine-treated mice (**Fig. S5C**). Histopathological assessment demonstrated that prolonged oral queuine administration preserved normal organ histology without significant damage. Notably, queuine treatment substantially improved structural integrity across multiple organs in aged mice (**Fig. S5E**).

### Queuine deficiency-induced reduction in manQ tRNA modification represents a novel epigenetic hallmark of aging

Cytosolic queuine is incorporated by tRNA-guanine transglycosylase (TGT) into the anticodon loop (position 34) of four tRNAs: tRNA^Asn^, tRNA^His^, tRNA^Asp^, and tRNA^Tyr.^ We therefore hypothesize that age-related queuine deficiency impairs queuosine biosynthesis and modification of target tRNAs. To map age-dependent changes in tRNA modifications, we isolated RNA from 2-, 6-, 24-, and 36-month-old rat tissues. Urea-PAGE confirmed RNA integrity (**Fig. 5A**). Analysis of 52 RNase T1-digested tRNA fragments via UHPLC-QTOF-MS (**Table S2**) revealed maximal fragmentation differences between 6- and 36-month-old rats by OPLS-DA (**Fig. 5B**). Specifically, the CUC[manQ]UCA[m^5^C]G fragment from tRNA^Asp^ was significantly downregulated in kidneys of 36-month-old versus 6-month-old rats (**Fig. 5C, D, F**), with cross-organ validation (**Fig. S6A**). Secondary MS confirmed fragment identity (**Fig. 5E**). Critically, the UAUCCCG fragment (same tRNA^Asp^ isoacceptor) showed no reduction (**Fig. 5F**), indicating modification-specific rather than transcript-level changes. Importantly, other queuosine-modified tRNA fragments remained unaltered (**Fig. 5C, D**), establishing manQ reduction as a selective age-associated event.

**Figure 5.**
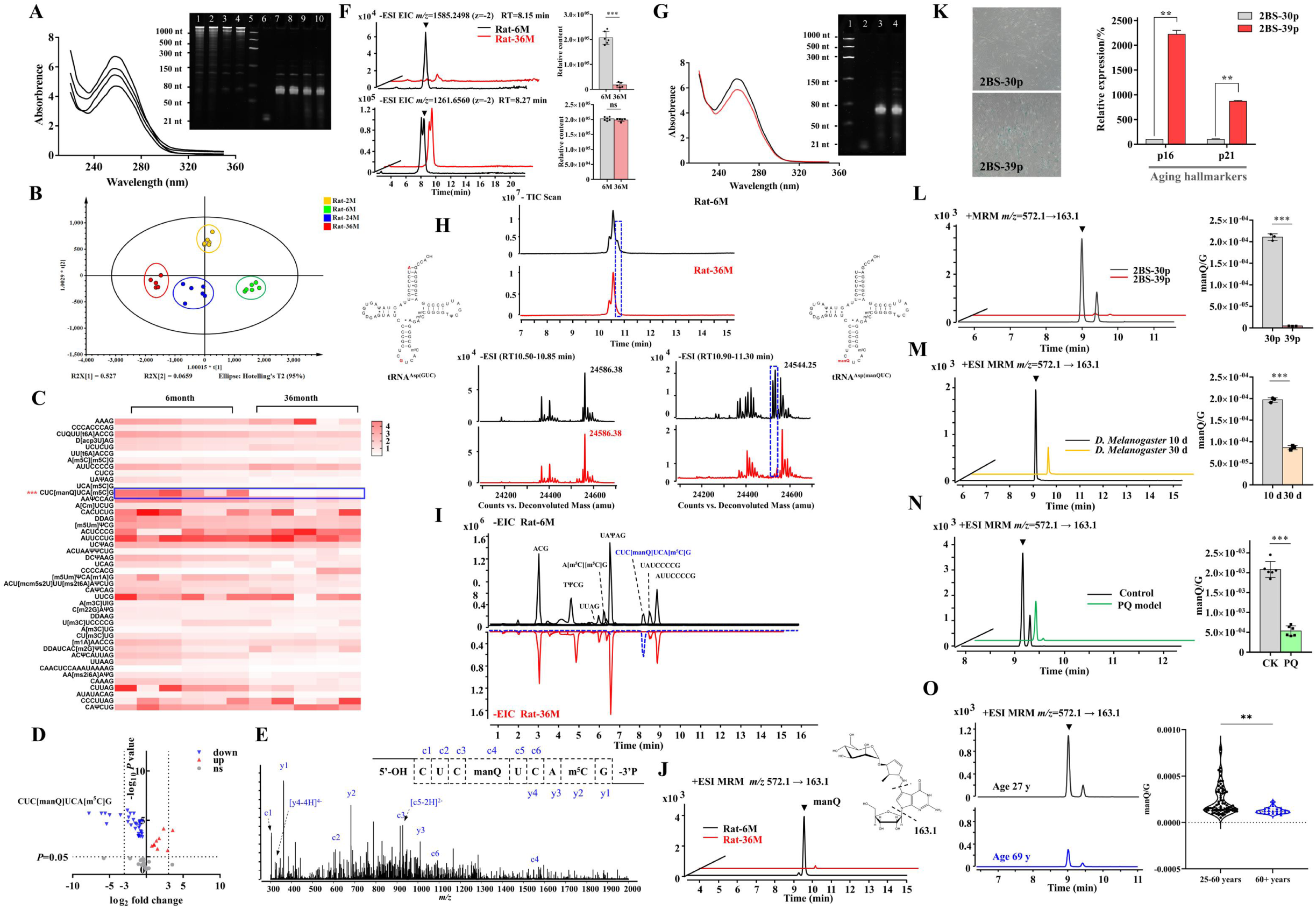
Queuine-dependent tRNA modification manQ deficiency emerges as a conserved epitranscriptomic hallmark of aging. (A) Quality control of rat kidney RNA isolation. Top: Urea-PAGE of large (lanes 1-4) and small RNA (lanes 7-10); Ladders (lane 5: Low range, lane 6: miRNA). Bottom: Small RNA UV spectra (Rat-6M, 2M, 36M, 24M). (B) OPLS-DA score plot of tRNA fragments across age groups (t[1] = 0.527, t[2] = 0.066). (C-D) Age-dependent tRNA fragment alterations: Heatmap (C) and volcano plot (D) comparing 6mo vs 36mo rats. (E) Structural validation of CUC[manQ]UCA[m^5^C]G fragment by UHPLC-QTOF-MS/MS. CID spectrum shows c/y-ion series (mass tolerance ±0.5 Da). (F) Top: EIC chromatograms and quantitation of CUC[manQ]UCA[m^5^C]G fragment (\*\*\**p*<0.001 vs 6mo). Bottom: Control fragment UAUCCCG (ns). All data normalized to TΨCG peak area. (G) Purification of tRNAᴬˢᵖ from rat kidneys. Left: Urea-PAGE (lane 1: Low range ladder; lane 2: miRNA ladder; lanes 3-4: tRNAᴬˢᵖ from 6mo/36mo rats). Right: UV spectra. (H) tRNAᴬˢᵖ characterization. Top: TIC chromatograms (6mo vs 36mo). Bottom: Deconvoluted MS spectra of tRNA^ᴬˢᵖ(GUC)^ and tRNA^ᴬˢᵖ(manQUC)^ with sequence mapping. (I) UHPLC-QTOF-MS profile of RNase T1-digested tRNAᴬˢᵖ fragments. (J) Nucleoside analysis of purified tRNAᴬˢᵖ by UHPLC-QQQ-MS. (K) Senescence phenotypes in tRNA modification-deficient 2BS cells: SA-*β*-gal staining (left) and senescence marker expression (right). (L-O) Conservation of manQ deficiency across models: 2BS cells (L), *D. melanogaster* (M), paraquat-induced aging mice (N), (O) Human leukocyte (\**p*<0.05, \*\**p*<0.01, \*\*\**p*<0.001 vs aging/controls).

To systematically characterize age-associated alterations in tRNA^Asp(manQUC)^, we performed single-tRNA purification from renal tissues of adult (6-month) and aged (24-month) rats. Urea-PAGE analysis confirmed the structural integrity of purified tRNAs from both age groups (**Fig. 5G**). High-resolution mass spectrometry revealed complete absence of tRNA^Asp(manQUC)^ (RT=10.90-11.30 min, MW=24,544.25) in aged rats, while unmodified tRNA^Asp(GUC)^ (RT=10.50-10.85 min, MW=24,586.38) remained unchanged (**Fig. 5H**). Consistently, RNase T1 digestion of purified tRNA^Asp^ yielded no CUC[manQ]UCA[m^5^C]G fragment in aged samples (**Fig. 5I**), and nucleoside analysis detected no manQ modification (**Fig. 5J, S6B**). These findings establish manQ hypomodification as the specific molecular defect underlying age-related tRNA^Asp(manQUC)^ deficiency. Intriguingly, parallel purification showed no age-dependent reduction in queuosine-modified tRNAs: tRNA^Asn(QUU)^, tRNA^His(QUG)^, or tRNA^Tyr(galQΨA)^ (**Fig. S6C-E**). This aligns with Thurm et al., who reported queuine incorporation hierarchy prioritizes tRNA^Asn^, tRNA^His^, and tRNA^Tyr^ in queuine-replete conditions, yet demonstrated tRNA^Asp^ exhibits the most rapid response to queuine scarcity (Thumbs et al., 2020).

Given the evolutionary conservation of aging hallmarks, we systematically validated manQ modification reduction across four model systems: naturally senescent human 2BS cells, *D. melanogaster*, paraquat-induced acutely aged mice, and aged human cohorts. Senescence in 39th-generation 2BS cells was confirmed by *β*-galactosidase staining and elevated p16/p21 expression (**Fig. 5K**). Using the characteristic manQ fragmentation ion (*m/z* 572.0→163.0, **Fig. S6B**), we observed significant manQ reduction in senescent 2BS cells, aged *D. melanogaster*, paraquat-treated mice (acute aging model) and elderly human subjects (**Fig. 5L-O**). These cross-species data establish queuine deficiency-mediated manQ loss as a conserved epitranscriptomic hallmark of aging.

### Queuine supplementation effectively restores manQ modification *in vitro*

Having localized age-dependent manQ reduction, we analyzed expression of key enzymes in its biosynthetic pathway across young/aged humans and rats (**Fig. 6A**). *GTDC1* expression was significantly downregulated in aged versus young subjects (human and rat) (**Fig. 6B**). Conversely, *Qtrt1* and *Qtrt2* expression increased with aging in both species. We propose that manQ deficiency triggers compensatory feedback upregulation of queuine-incorporating enzymes (*Qtrt1/Qtrt2*) to maximize queuine utilization. This model is supported by maintained abundance of queuosine-modified tRNAs (tRNA^Asn(QUU)^, tRNA^His(QUG)^, tRNA^Tyr(galQΨA)^) in aged rats despite *GTDC1* downregulation (**Fig. S6C-E**).

**Figure 6.**
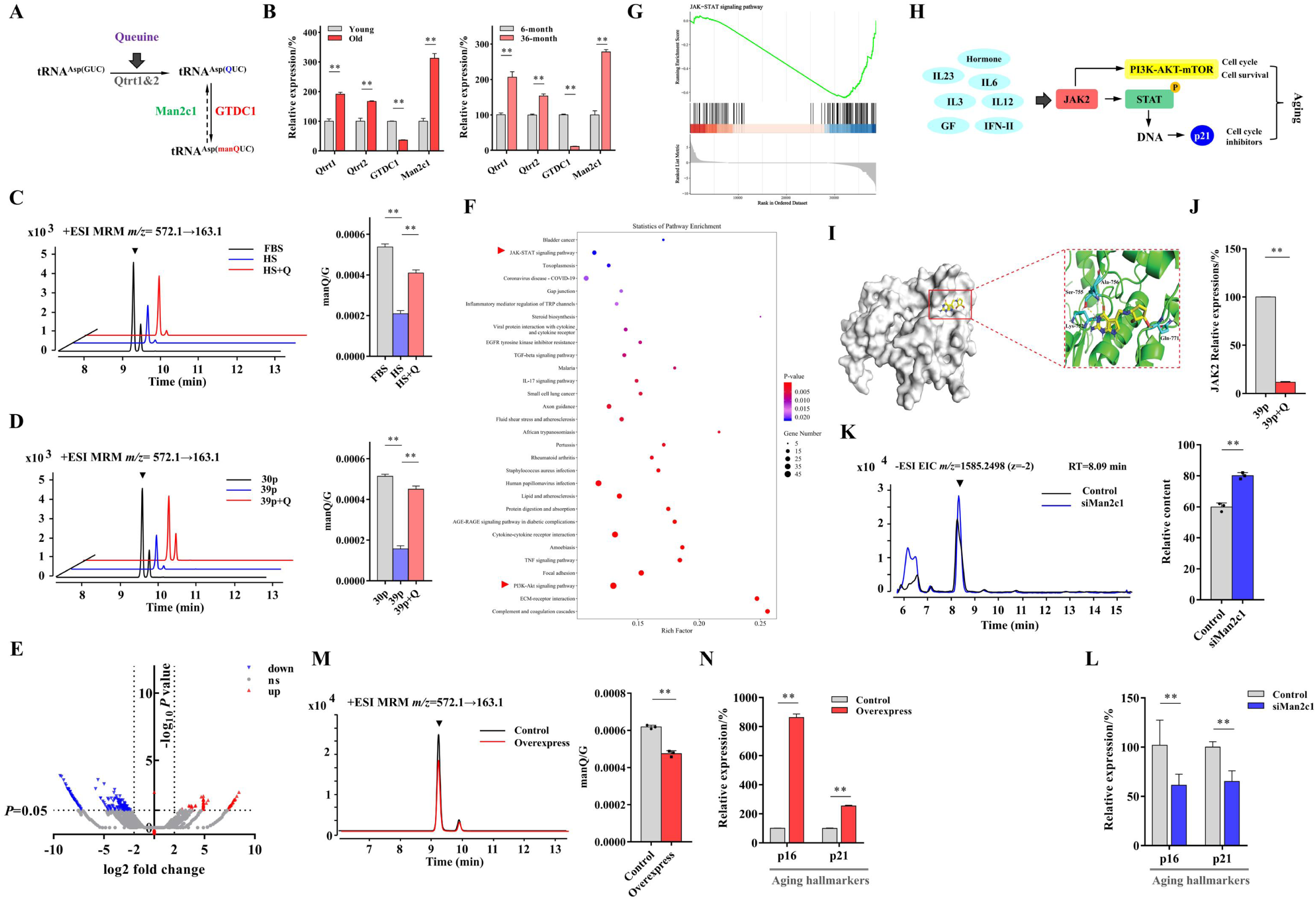
Targeted manipulation of manQ tRNA modification regulates cellular senescence (A) Biosynthetic pathway of manQ tRNA modification in humans. (B) Age-dependent dysregulation of manQ pathway enzymes: Left: Relative expression analysis in human leukocytes (young vs aged donors). Right: qPCR validation in rat kidneys (6mo vs 36mo) (C-D) UHPLC-MS quantification of manQ modification: (C) 2BS cells cultured in queuine-supplemented horse serum. (D) Naturally senescent 2BS cells (\**p*<0.05, \*\**p*<0.01). (E-G) Transcriptomic profiling of queuine-treated senescent 2BS cells: (E) Volcano plot of differentially expressed genes (|log₂FC|>2, FDR<0.05). (F) Top enriched KEGG pathways (bubble size: gene count; color: -log₁₀[p-value]). (G) GSEA showing JAK-STAT pathway suppression (NES=-2.1, FDR=0.002) (H) Human JAK-STAT signaling schematic highlighting queuine’s putative targeting. (I) Molecular docking of queuine in JAK2 kinase domain (binding affinity: −4.61 kcal/mol). (J) RT-qPCR showing JAK2 downregulation in queuine-treated 2BS cells. (K-L) *MAN2C1* knockdown consequences: (K) manQ reduction by UHPLC-MS (\*\**p*<0.01). (L) Senescence marker induction (p16/p21). (M-N) *MAN2C1* overexpression effects: (M) manQ restoration (\*\**p*<0.01 vs control). (N) Senescence marker suppression (\*\**p*<0.01).

Supplementation with queuine (10 ng/mL) in horse serum-cultured senescent 2BS cells elevated manQ modification ∼2-fold versus controls *in vitro* (**Fig. 6C,D**), consistent with Thurm et al. RNA-seq analysis revealed significantly reduced JAK2 expression in queuine-treated versus untreated senescent cells (**Fig. 6E**). KEGG pathway analysis confirmed inhibition of JAK-STAT and PI3K-Akt signaling in treated cells (**Fig. 6F,G,S7B**). GO enrichment indicated altered DNA-binding transcription repressor activity (RNA polymerase II-specific) in queuine-treated cells (**Fig. S7A**). Mechanistically, aging-associated inflammation (e.g., IL-6 upregulation) activates JAK2-STAT signaling to modulate transcription (indirectly regulating p21) and crosstalks with PI3K/AKT/mTOR pathways to control proliferation, differentiation, and apoptosis (**Fig. 6H**). Notably, molecular docking demonstrated high-affinity queuine-JAK2 binding (docking energy: −4.61 kJ/mol) (**Fig. 6I**), indicating strong hydrogen bonding interactions with amino acids (Ala-756, Ser-755, Lys-752, Gln-771). Cell experiments further confirmed that queuine suppressed JAK2 expression (**Fig. 6J**), functionally decoupling its epigenetic effects from kinase regulation.

Furthermore, we systematically profiled mannose hydrolases and focused on Man2c1, an enzyme mediating free oligosaccharide (fOSs) metabolism by cleaving mannose from misfolded glycoprotein side chains, as dysregulated in aging (Maia et al., 2022). Paradoxically, *MAN2C1* expression significantly increased in aged humans and rats versus young controls (**Fig. 6B**). We hypothesized that aberrant Man2c1 accumulation promiscuously degrades manQ due to substrate non-specificity. siRNA-mediated *MAN2C1* knockdown rescued manQ modification levels in 2BS cells (**Fig. 6K**) and suppressed senescence markers p16/p21 (**Fig. 6L**). *MAN2C1* overexpression reduced manQ modification (**Fig. 6M**) while elevating p16/p21 expression (**Fig. 6N**) demonstrating Man2c1 as a functionally conserved regulator of manQ-dependent senescence.

### Reduced manQ modifications disrupt proteostasis via impaired translation

CRISPR-Cas9-mediated targeting of *GTDC1* (**Fig. 7A**) achieved >50% knockdown efficiency (**Fig. 7B**), significantly reducing manQ levels (**Fig. 7C**) and accelerating senescence in 2BS cells (**Fig. 7D,E**). DIA quantitative proteomics revealed widespread proteomic dysregulation in *GTDC1*-KD cells (>1,000 aberrantly expressed proteins; **Fig. 7F-H**). Key findings include: senescence-associated kinases upregulation (PDGFRA, MAPK7, AXL, CDK12; **Fig. 7L**), GPNMB downregulation (novel aging marker regulating mitophagy via mTOR) (**Fig. 7H,I**) (Yu et al., 2024), Man2c1 upregulation (validating **Fig. 6B**) consistent (**Fig. 7K**). KEGG/GO analyses implicated ribosomal dysfunction and translation metabolism pathways (**Fig. S8A,B**). Strikingly, differentially expressed proteins were enriched in neurodegenerative pathways (Alzheimer’s, Huntington’s, Parkinson’s), mechanistically linking manQ deficiency to proteostatic collapse (**Fig. 7J**).

**Figure 7.**
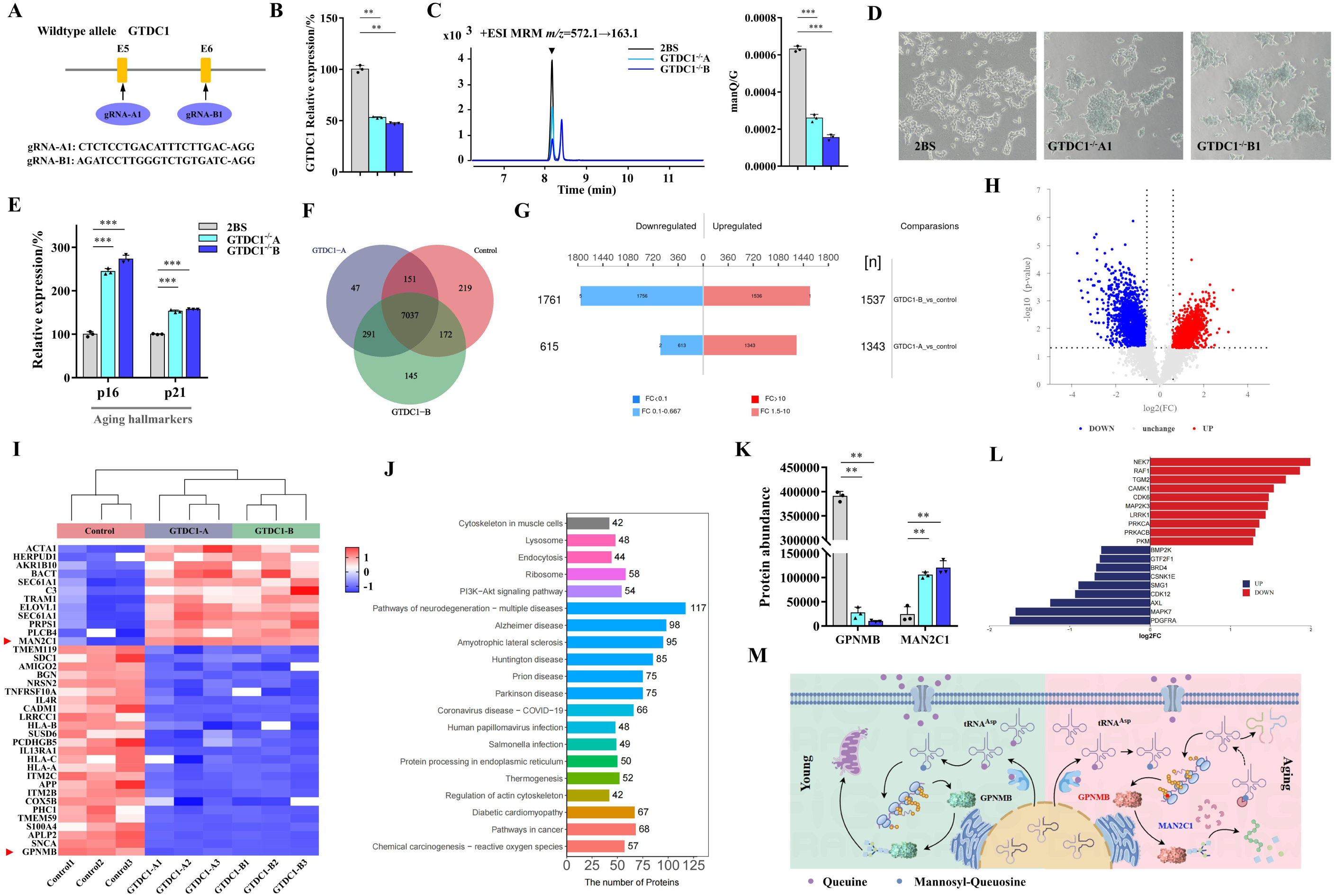
Aging characterized by loss of proteostasis is associated with reduced manQ modification. (A) CRISPR/Cas9-mediated *GTDC1* knockout strategy in 2BS cells. (B) Validation of GTDC1 ablation efficiency by qPCR. (C-E) Consequences of manQ loss: (C) UHPLC-MS quantification of manQ modification (\*\*\**p*<0.001 vs WT). (D) Increased SA-*β*-gal+ cells. (E) Senescence marker upregulation (p16/p21; \*\*\**p*<0.001) (F-L) Proteomics of WT vs *GTDC1*^-/-^ 2BS cells: (F) Shared/unique proteins (Venn diagram). (G) Differential protein abundance histogram (FDR<0.05). (H) Volcano plot of dysregulated proteins. (I) Hierarchical clustering of altered proteins. (J) Significant differential protein KEGG pathway annotation and attribution histogram (K) Validation of GPNMB depletion and Man2c1 accumulation (\*\**p*<0.01 vs WT). (L) Kinase activity landscape (top 10 upregulated/downregulated). (M) Mechanistic model: manQ deficiency impairs tRNA decoding fidelity-ribosomal stalling-proteotoxic stress-senescence via GPNMB/MAPK activation.

Mechanistically, queuine uptake in young organisms sustains physiological manQ modification levels, ensuring translational fidelity of key proteins (e.g., GPNMB) and proteostatic homeostasis, including mitochondrial autophagy. With aging, however, queuine insufficiency and *GTDC1* downregulation impair manQ synthesis, driving aberrant expression of GPNMB and other regulators. Concurrent Man2c1 upregulation, while facilitating clearance of misglycosylated proteins, promiscuously degrades manQ due to substrate promiscuity (**Fig. 7M**). Thus, manQ deficiency initiates a paradoxical cycle: as both a driver amplifying proteostatic collapse and a consequence of age-related metabolic dysfunction, it epitomizes the self-reinforcing nature of epigenetic aging.

### Queuine supplementation demonstrates systemic metabolic reprogramming capacity *in vivo*

Untargeted metabolomics of mouse plasma (young, aged, queuine-treated aged; n=539 metabolites) revealed significant metabolic reprogramming in queuine-treated aged mice versus controls (**Fig. 8A-D**). Anti-inflammatory prostaglandins (6-keto-PGF1α) and specialized pro-resolving mediators (11-HEPE, 5-HEPE) were elevated (**Fig. 8E**). Kynurenine pathway was attenuated (delaying neurodegeneration) (**Fig. 8F**). Increased nicotinamide ribotide (validated anti-aging metabolite) was also observed (**Fig. 8F**). These coordinated shifts position queuine as a metabolic regulator that counteracts age-related biochemical dysfunction.

**Figure 8.**
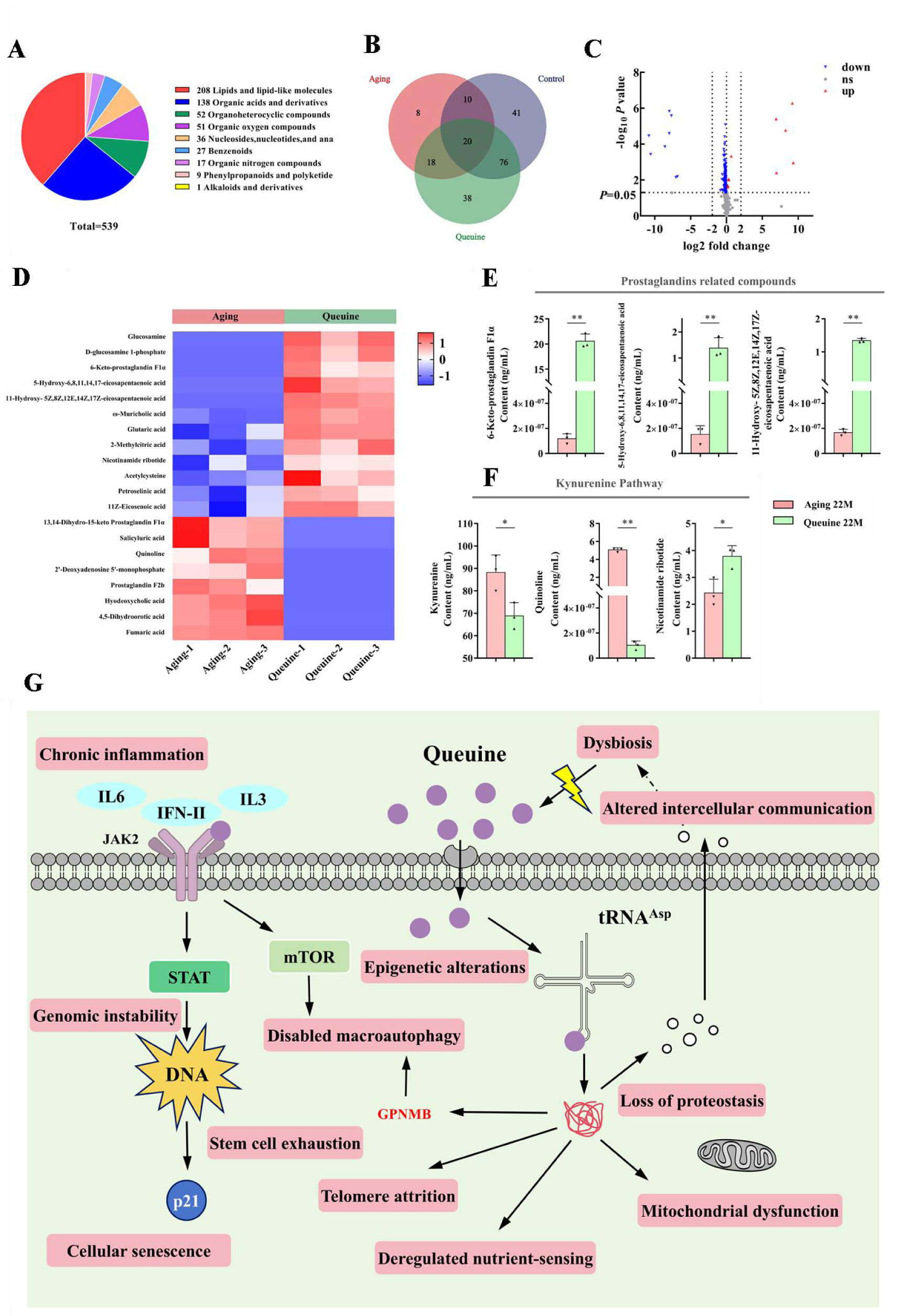
Systemic queuine supplementation counteracts metabolic dysregulation in aging. (A-B) Plasma metabolomics profiling: (A) Proportional distribution of metabolite classes, (B) Shared/unique metabolites across groups (young/aged/aged+queuine). (C-D) Dysregulated metabolites in aged vs. queuine-treated mice: (C) Volcano plot (|log_2_FC|>2, FDR<0.05), (D) Hierarchical clustering of altered metabolites. (E-F) Pathway-specific metabolite restoration: (E) Prostaglandins, (F) Kynurenine pathway (\**p*<0.05,\*\**p*<0.01). (G) Integrative model: Queuine-mediated manQ restoration modulates 12 hallmarks of aging.

The regulatory role of queuine in aging involves multi-layered mechanisms. Age-related queuine deficiency, driven by gut dysbiosis and chronic inflammation, compromises cellular translocation efficiency, impairing tRNA manQ synthesis. This reduces translational fidelity, disrupts proteostasis, and crosstalks with classical aging hallmarks, such as nutrient sensing downregulation, mitochondrial dysfunction, altered intercellular communication and Telomere attrition. Mechanistically, queuine competitively inhibits inflammatory cytokine receptors (e.g., JAK2), blocking IL-6/IFN-γ/IL-3 signaling to suppress STAT/mTOR-mediated DNA damage, senescence, and mitophagy impairment. Simultaneously, it restores manQ-dependent translational accuracy, ensuring precise expression of senescence regulators like GPNMB. Consequently, mitochondrial function, nutrient sensing, telomerase activity, and intercellular communication are progressively rescued through proteostatic remodeling. Critically, queuine establishes a positive feedback loop: improved host physiology reverses gut dysbiosis, enriching queuine-producing symbionts to amplify anti-aging effects. This multi-targeted action collectively reprograms twelve hallmarks of aging (**Fig. 8G**).

## DISCUSSION

Aging is increasingly viewed as a systems-level process driven by interconnected molecular failures, yet the upstream regulatory events that coordinate these hallmarks remain incompletely defined. Although the hallmarks framework provides a powerful conceptual scaffold, many hallmarks primarily describe downstream cellular and tissue consequences rather than the primary molecular defects that initiate or synchronize functional decline. Here, we identify an evolutionarily conserved, age-dependent depletion of tRNA mannosyl-queuosine (manQ) as a previously underappreciated epitranscriptomic alteration that links translational control to organismal aging. By integrating multi-organ tRNA modification profiling with targeted genetic perturbations and nutritional restoration across species, our data position manQ decline at a critical regulatory nexus connecting decoding fidelity, proteome integrity, and systemic deterioration.

Among the more than one hundred known RNA modifications, tRNA modifications constitute the most abundant and structurally diverse class, yet their contributions to aging biology remain comparatively understudied. Our profiling establishes manQ as a uniquely consistent tRNA-specific modification that declines across phylogenetically distant models and diverse senescence paradigms. Importantly, manQ hypomodification is selective rather than reflecting global tRNA depletion: aging preferentially reduces the manQ-containing tRNA^Asp^ fragment while leaving the corresponding unmodified tRNA^Asp^ fragment, and other queuosine-modified tRNAs relatively unchanged. This pattern supports a regulated defect in modification homeostasis rather than a generalized change in transcript abundance. Such specificity argues that manQ loss is not merely a passive consequence of tissue degeneration, but instead represents a conserved, biologically meaningful aging-associated event with mechanistic impact.

A central conceptual advance of our work is the positioning of manQ as a functionally informative biomarker candidate that aligns with widely accepted criteria for robust aging indicators: (i) a quantifiable trajectory across age, (ii) causal linkage to aging phenotypes, and (iii) therapeutic reversibility upon targeted intervention. First, we observe an age-associated decline in circulating queuine — the microbiota-derived precursor required for manQ biosynthesis — in both rodents and humans, consistent with a systemic determinant that can be measured reproducibly across aging. Second, experimental depletion of manQ through pathway perturbation accelerates senescence programs, whereas restoration of queuine availability replenishes manQ and suppresses p16/p21-linked senescence signatures across cellular and organismal contexts. Third, queuine supplementation confers therapeutic reversibility across biological scales, improving healthspan phenotypes and extending lifespan in Drosophila and naturally aging mice. Together, these findings elevate manQ from a correlative molecular signature to a mechanistically actionable feature of biological aging.

Mechanistically, manQ occupies a privileged position in gene expression: the wobble nucleotide (position 34) of tRNA^Asp^, where codon–anticodon pairing constraints and decoding kinetics are established. This location provides a direct route by which age-associated manQ loss can propagate broadly through the proteome. Our data support a model in which manQ functions as a safeguard of translational fidelity, and its depletion compromises decoding accuracy and/or elongation dynamics, increasing the burden of aberrant polypeptides and destabilizing proteome balance. Consistent with this framework, manQ deficiency is associated with widespread proteomic remodeling and altered abundance of aging-relevant proteins, suggesting a proteostasis-centered mechanism whereby translational infidelity becomes an upstream driver of stress response activation, inflammatory signaling, and reinforcement of senescence programs. Because proteostasis intersects with multiple canonical hallmarks (e.g. mitochondrial dysfunction, impaired stress resilience, and altered intercellular communication), translation-coupled proteome destabilization offers a unifying explanation for how a single tRNA modification defect can elicit multi-system consequences. In this view, manQ decline is not merely one of many molecular changes observed in aging, but rather a proximate determinant capable of amplifying downstream hallmarks through a common axis of proteome quality control.

Our findings further suggest that manQ depletion may engage self-reinforcing feedback loops that accelerate aging trajectories. Aging is accompanied by reduced queuine availability and additional systemic constraints, including inflammatory states that may impair queuine uptake and utilization, thereby further limiting manQ biosynthesis. At the same time, the selective nature of manQ depletion—particularly the sensitivity of tRNA^Asp^ to queuine scarcity compared with other queuosine-modified tRNAs—may render translational decoding vulnerable even under modest reductions in circulating queuine. This architecture offers a conceptual framework in which aging progressively erodes “epitranscriptomic integrity” at the tRNA level, pushing translation toward an error-prone regime that accelerates proteostatic collapse and functional decline. Such a model also provides a mechanistic rationale for the breadth of phenotypic rescue observed upon queuine repletion: restoring a proximal fidelity determinant can reduce proteotoxic burden and thereby dampen multiple downstream stress and senescence pathways simultaneously.

A distinctive implication of this work is that queuine introduces a microbiota–host epitranscriptomic axis into aging biology. Queuine is produced by gut microbiota and cannot be synthesized de novo by mammals, making it an environmentally and nutritionally contingent determinant of host tRNA modification state. The observation that circulating queuine declines with age in rodents and humans suggests that age-associated microbial and/or host physiological changes can translate into a conserved deficit in a tRNA modification precursor. Notably, long-term queuine supplementation in mice is accompanied by remodeling of gut microbial community structure and metabolic outputs, including increased abundance of putatively beneficial taxa and elevated short-chain fatty acids. These patterns are consistent with bidirectional reinforcement between host physiology and microbial ecology and suggest that queuine may act not only as a passive microbial product but also as a modulator of host–microbiome homeostasis. Collectively, these findings expand the conceptual scope of geroscience by placing a microbiota-derived nutrient upstream of translational quality control and by suggesting that age-associated dysbiosis may exert part of its influence through perturbation of host epitranscriptomic regulation.

From a translational perspective, most candidate geroprotective interventions target nutrient-sensing pathways, autophagy, or senescent-cell clearance. While these approaches have yielded important mechanistic insights, each typically engages only a subset of aging mechanisms and often raise concerns about durability, heterogeneity of response, or long-term tolerability. Queuine supplementation offers a distinct therapeutic logic: rather than modulating a single signaling cascade, it restores a tRNA modification state that governs translational fidelity—an upstream determinant of proteome quality that can, in principle, influence multiple downstream hallmarks concurrently. In line with this systems-level mechanism, queuine extends median lifespan by ∼47% in Drosophila and increases mean lifespan by ∼15.3% in naturally aging mice, accompanied by broad improvements in functional measures. Although cross-species effect sizes should be interpreted within the context of model-specific biology, the consistency of benefit across evolutionarily distant organisms supports the premise that maintaining translational accuracy via tRNA epitranscriptomic integrity may represent a multi-target route to geroprotection. More broadly, these findings highlight an intervention paradigm centered on restoring molecular fidelity, rather than suppressing a single downstream phenotype, as a strategy to delay systemic aging.

Several limitations and open questions should guide future work. First, the precise spectrum of codon-specific decoding errors and kinetic alterations induced by manQ depletion remains to be fully defined. Integrating ribosome profiling with quantitative proteomics that directly measures amino acid misincorporation would strengthen the mechanistic chain from manQ status to proteome damage. Second, the drivers of age-associated queuine decline are likely multifactorial, potentially involving shifts in microbiota composition, altered intestinal absorption, and inflammation-linked inhibition of uptake or utilization pathways. Dissecting these contributions will be essential for developing precision interventions and for identifying populations most likely to benefit. Third, although our long-term mouse studies did not detect overt toxicity, translation-layer interventions could, in principle, carry trade-offs that manifest only over extended periods or in specific physiological contexts. Rigorous pharmacokinetic profiling, tissue distribution, and long-term safety evaluations, together with assessment of potential impacts on proteome diversity and stress adaptability, will be critical prerequisites for clinical translation.

In summary, our study identifies a conserved, age-associated loss of tRNA mannosyl-queuosine as an epitranscriptomic signature that links microbiota-derived nutrient availability to translational fidelity and proteome integrity. Selective manQ depletion in aging provides a mechanistically interpretable route from erosion of tRNA modification homeostasis to proteostasis collapse and multi-hallmark functional decline. Restoration of manQ through queuine supplementation improves translational control and confers broad benefits across species, establishing tRNA epitranscriptomic remodeling as a tractable and potentially actionable layer of systemic aging regulation. More generally, our findings advance a conceptual framework in which maintenance of translational accuracy at the tRNA level constitutes a foundational determinant of aging trajectories and a promising target for interventions aimed at extending healthspan.

## ACKNOWLEDGMENTS

We acknowledge Figdraw (www.figdraw.com) for graphic illustration support. Technical assistance was provided by: Guangzhou GeneCopoeia (stable cell line construction), Guangzhou Saiye Biotech (gene knockout cell lines), Guangzhou Ruibo Biotech (transcriptome sequencing), BGI Genomics (DNA primer synthesis), Suzhou Beixin Biotechnology (siRNA synthesis), Shanghai Zhongke New Life (proteomics and targeted metabolomics), Shanghai Pisenor (16S rRNA sequencing and SCFA analysis), Shenzhen Aisi Gene (DNA methylation clock analysis).

## FOUNDATION

This work was financially supported by Macao Science and Technology Development Fund, Macau SAR (File no. 006/2023/SKL, 0001/2023/AKP) (Z.H.J.). This work was partially supported by Ruina (Zhuhai Hengqin) Biothechnology.

## SUPPLEMENTAL INFORMATION

Supplemental Figure 1. Queuine-dependent tRNA modifications in human cells.

Supplemental Figure 2. Queuine is significantly depleted in horse serum.

Supplemental Figure 4. Queuine remodels gut microbiota diversity and metabolic output in naturally aging male mice.

Supplemental Figure 5. Queuine treatment modulates other age-related health parameters in naturally aged mice.

Supplemental Figure 6. Tissue-specific profiling of queuine-modified tRNAs reveals selective age-dependent changes.

Supplemental Figure 7. Transcriptomic landscape of queuine-mediated senescence regulation.

Supplemental Figure 8. Functional annotation of *GTDC1* knockout-induced proteome dysregulation.

Supplemental Table 1 UHPLC-QQQ-MS instrument parameters of plasma metabolites.

Supplemental Table 2 Detailed mass spectrometry information of 52 tRNA fragments from rats using UHPLC-QTOF-MS.

