## Supplemental Figure1-8,and Supplemental Table 1-2 for "Evolutionarily Conserved Decline of tRNA Mannosyl-Queuosine Links Translational Regulation to Aging and Is Reversed by Queuine"

|  | 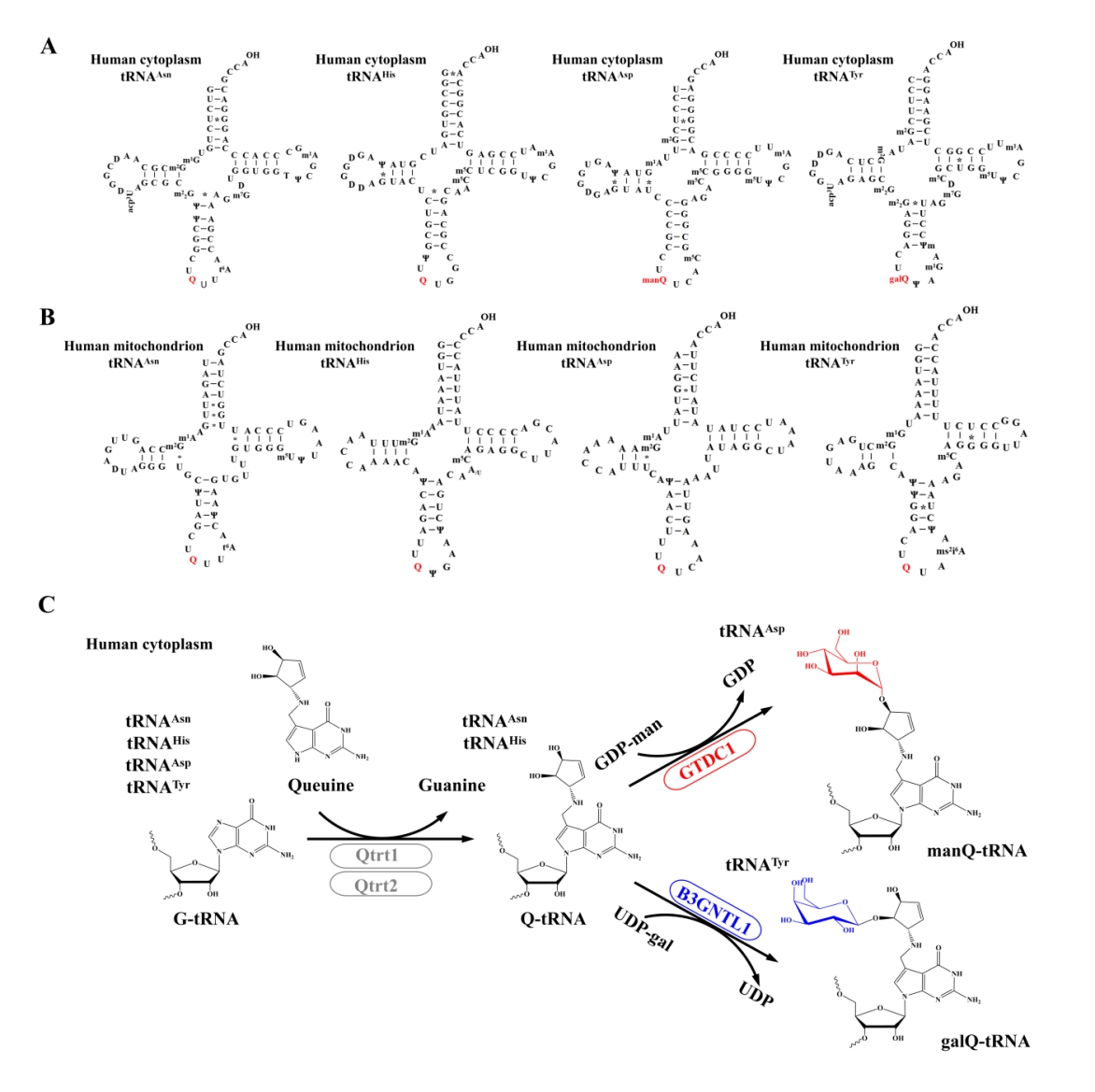 |
| --- | --- |

**Supplemental Figure 1. Queuine-dependent tRNA modifications in human cells.**

(A) Cytoplasmic tRNA substrates of queuine modification. Secondary structures of the four human cytoplasmic tRNAs incorporating queuine-derived modifications at position 34: queuosine (Q) in tRNA^Asn^ and tRNA^His^, mannosyl-queuosine (manQ) in tRNA^Asp^, and galactosyl-queuosine (galQ) in tRNA^Tyr^.
(B) Mitochondrial tRNA substrates of queuine modification. Secondary structures of the four human mitochondrial tRNAs modified by queuosine (Q) at position 34: tRNA^Asn^, tRNA^His^, tRNA^Asp^, and tRNA^Tyr^.
(C) Queuine modification pathway in human cytoplasm. Biosynthetic pathway of queuosine and its glycosylated derivatives (manQ, galQ) in human cells.

|  | 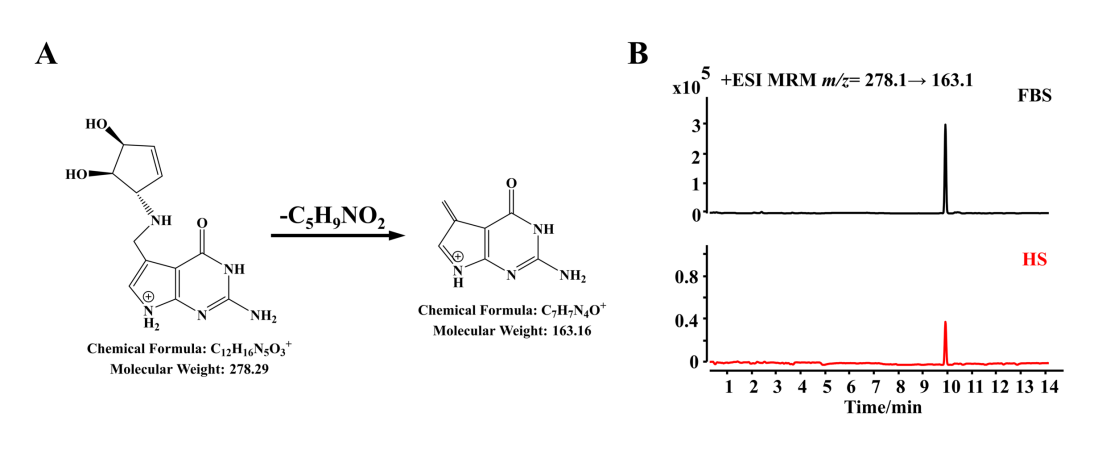 |
| --- | --- |

**Supplemental Figure 2. Queuine is significantly depleted in horse serum.**

(A) Representative fragmentation pattern of queuine standard (precursor ion: *m/z*=278.29 [M+H]⁺; characteristic product ion: *m/z=*163.16).

(B) Representative UHPLC-QQQ-MS chromatograms showing queuine peaks (retention time: 9.9 min) in fetal bovine serum (FBS) vs. horse serum (HS), demonstrating >8.3-fold reduction in HS (peak area ratio: HS/FBS = 0.12).

|  | 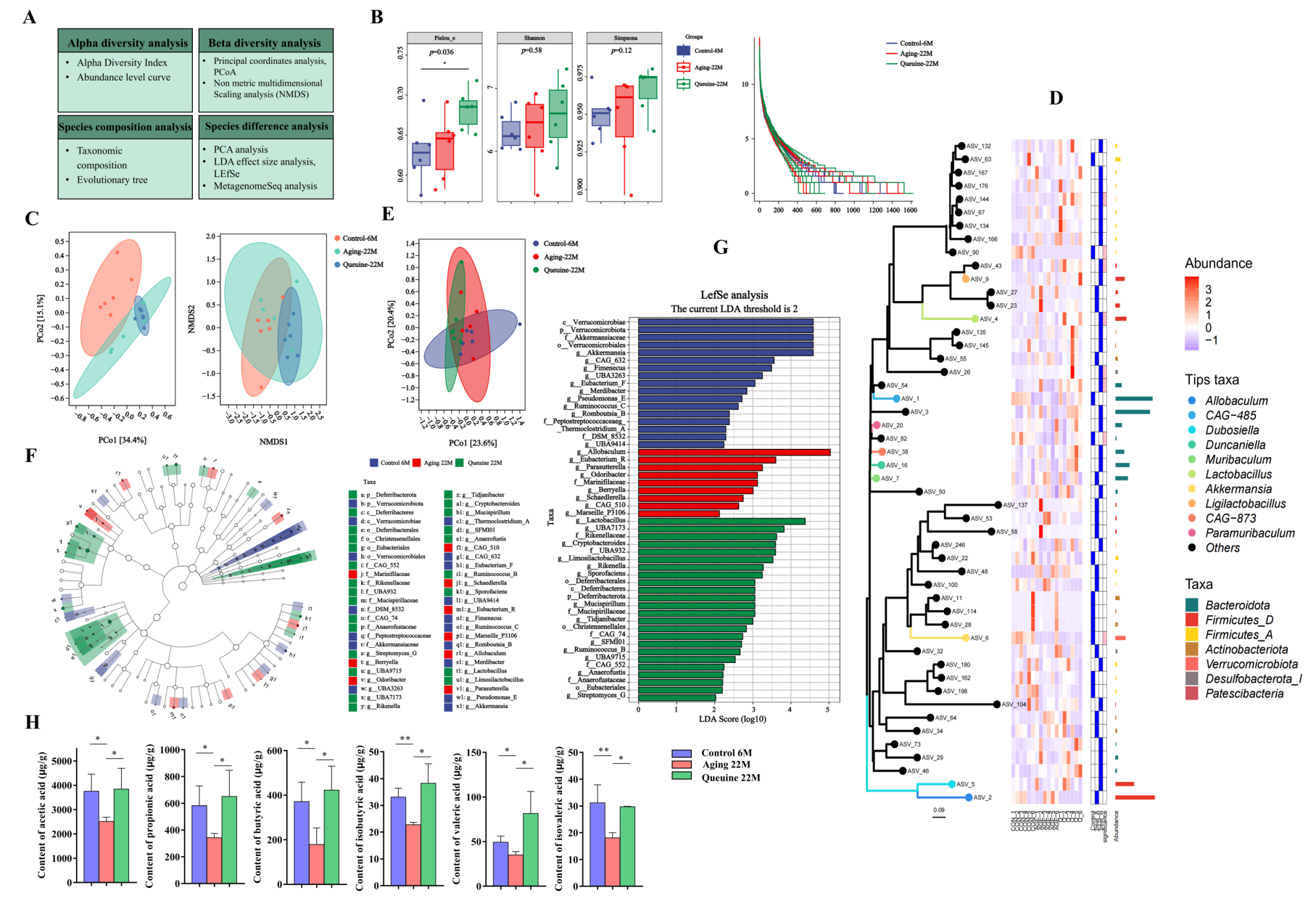 |
| --- | --- |

**Supplemental Figure 4. Queuine remodels gut microbiota diversity and metabolic output in naturally aging male mice.**

(A) Experimental design for gut microbiota and short-chain fatty acid (SCFA) analysis in young (6mo), aged (22mo), and aged+queuine (22mo) male C57BL/6J mice (n=6/group).

(B-G) Microbial community analysis: α-Diversity: Pielou's evenness, Shannon index, Simpson index, and rank-abundance curves (B). β-Diversity: Principal Coordinates Analysis (PCoA; Bray-Curtis) and Non-Metric Multidimensional Scaling (NMDS) (C). Phylogenetic structure: Maximum-likelihood phylogenetic tree (D). Compositional variance: Principal Component Analysis (PCA) of species abundance (E). Taxonomic biomarkers: LEfSe analysis showing differentially abundant taxa (branching diagram (F); LDA score distribution (G))

(H) Queuine-induced alterations in cecal short-chain fatty acid (SCFA) profiles. Concentrations of acetate, butyrate, isobutyrate, isovalerate, propionate, and valerate measured by GC-MS. Data presented as mean ± SEM; **p*<0.05, ***p*<0.01 vs. aged controls (two-way ANOVA with Tukey's post-hoc).

|  | 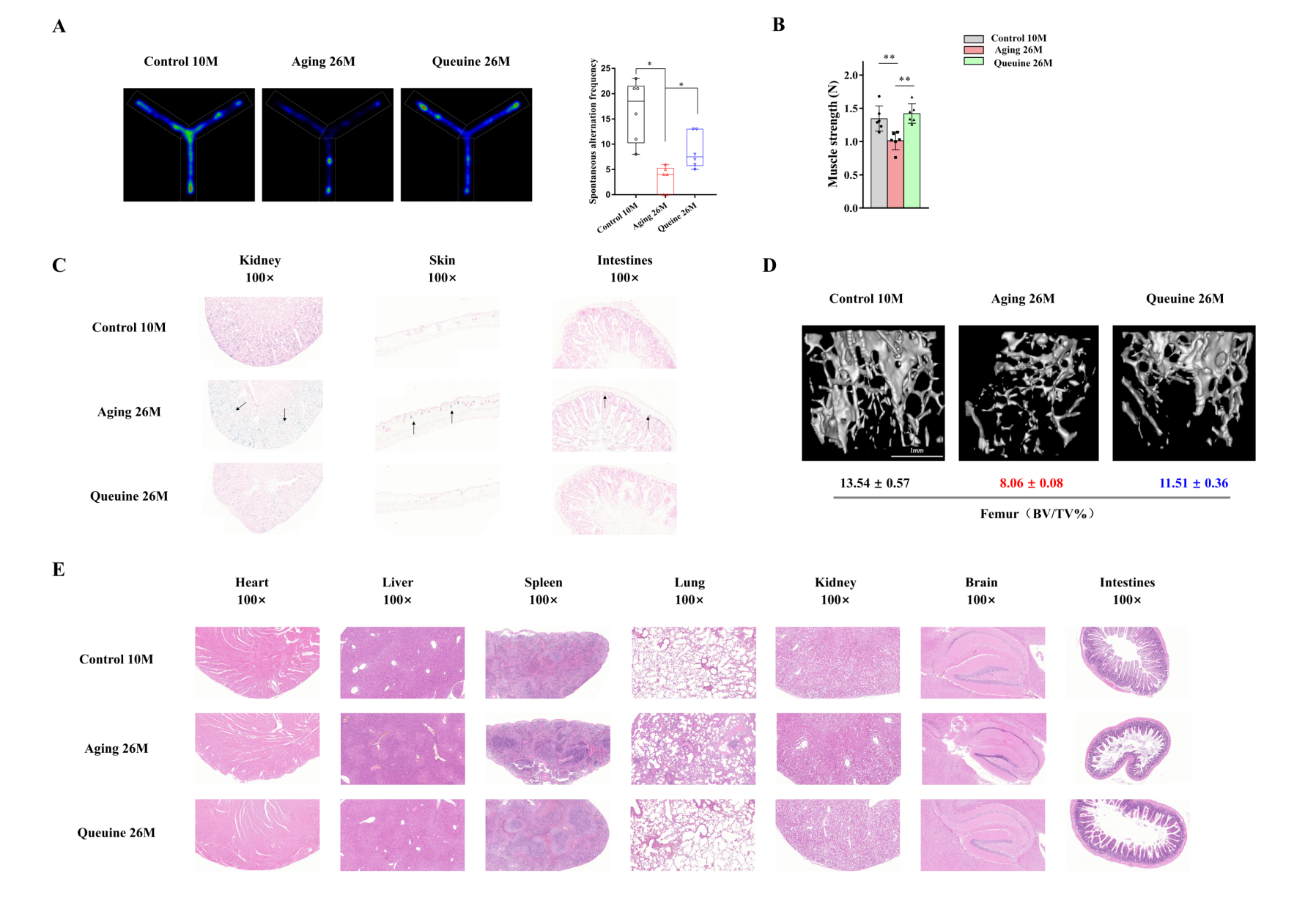 |
| --- | --- |

**Supplemental Figure 5. Queuine treatment modulates other age-related health parameters in naturally aged mice.**

(A-D) Effects of queuine administration on cognitive performance (Y-maze), muscle function (grip strength), cellular senescence (β-galactosidase activity), and skeletal integrity (bone mineral density) in 24-month-old C57BL/6 mice.

(E) Histopathological analysis of major organ systems following chronic queuine supplementation (12-month treatment) in aged cohorts.

|  | 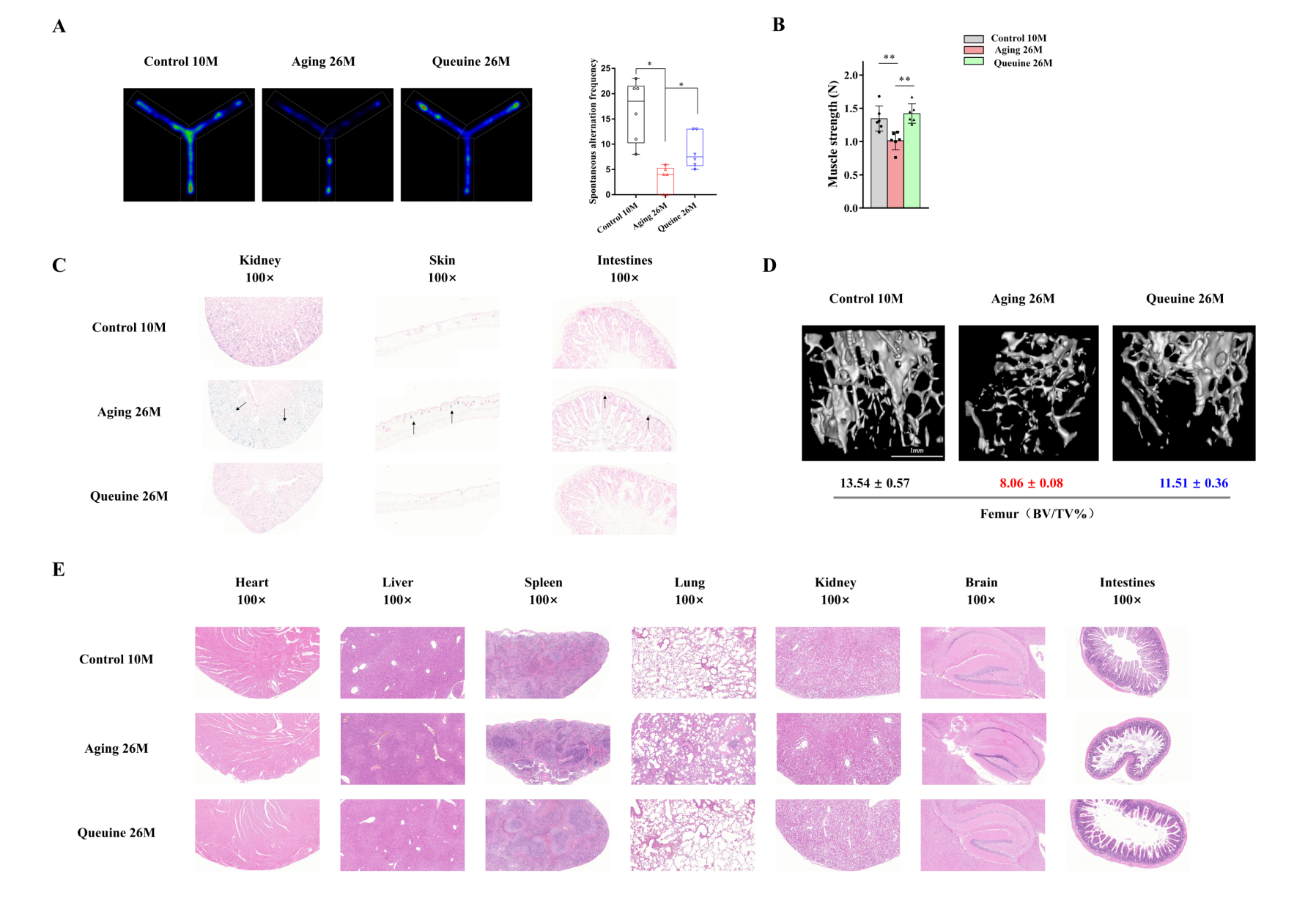 |
| --- | --- |

**Supplemental Figure 6. Tissue-specific profiling of queuine-modified tRNAs reveals selective age-dependent changes.**

(A) Quantitative UHPLC-MS/MS analysis of CUC[manQ]UCA[m^5^C]G fragment across multiple organs in 6-month-old (adult) versus 36-month-old (aged) rats. Data normalized to internal standard, **p*<0.05 , ***p*<0.01，****p*<0.001 vs. adult (two-way ANOVA).

(B) Characteristic MS/MS fragmentation pattern of synthetic mannosyl-queuosine (manQ) standard. CID spectrum shows diagnostic ions at *m/z* 410.41, 295.27, 163.16 (collision energy: 20 eV).

(C-E) Stability analysis of queuine-modified tRNAs in aged rat kidneys: tRNA^Asn(QUU)^ modification status (C), tRNA^His(QUG)^ relative abundance (D), tRNA^Tyr(galQΨA)^ modification profile (E). No significant alterations observed between adult (6mo) and aged (36mo) rats (ns *p*>0.05).

|  | 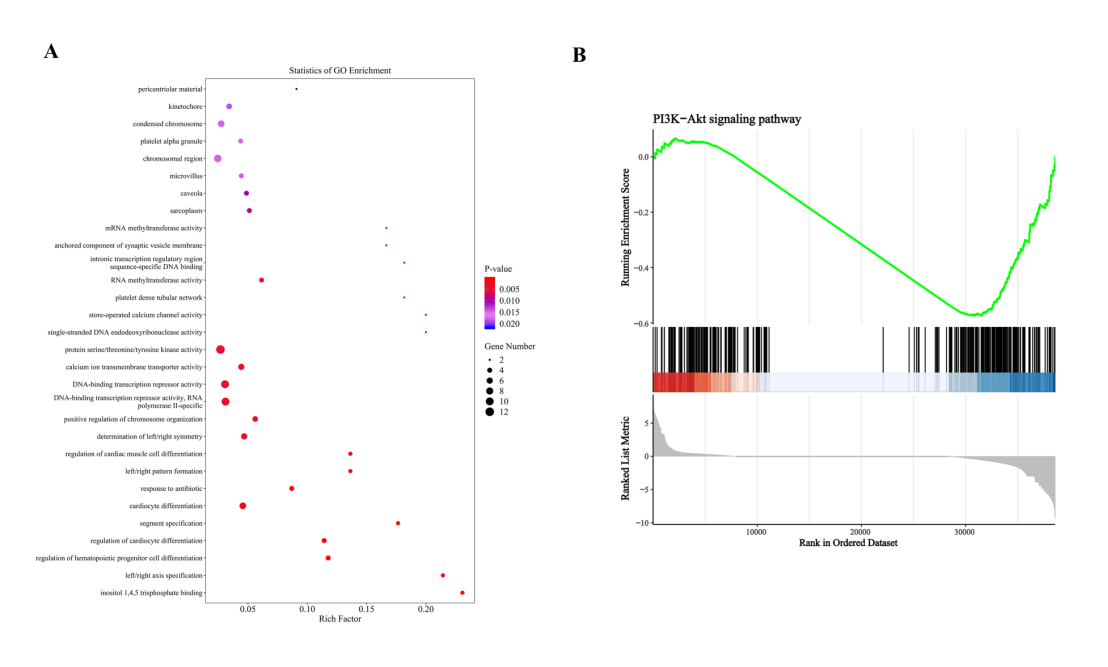 |
| --- | --- |

**Supplemental Figure 7. Transcriptomic landscape of queuine-mediated senescence regulation.**

(A) Gene Ontology (GO) enrichment of differentially expressed genes (|log_2_FC|>2, FDR<0.05) between naturally senescent and queuine-supplemented 2BS cells. Bubble size indicates gene count; color scale reflects -log_10_(FDR) significance.

(B) GSEA confirmation of PI3K-Akt pathway suppression (Normalized Enrichment Score [NES] = -1.87, FDR = 0.023). Gene rank metric shown with leading edge subset (black tick marks).

|  | 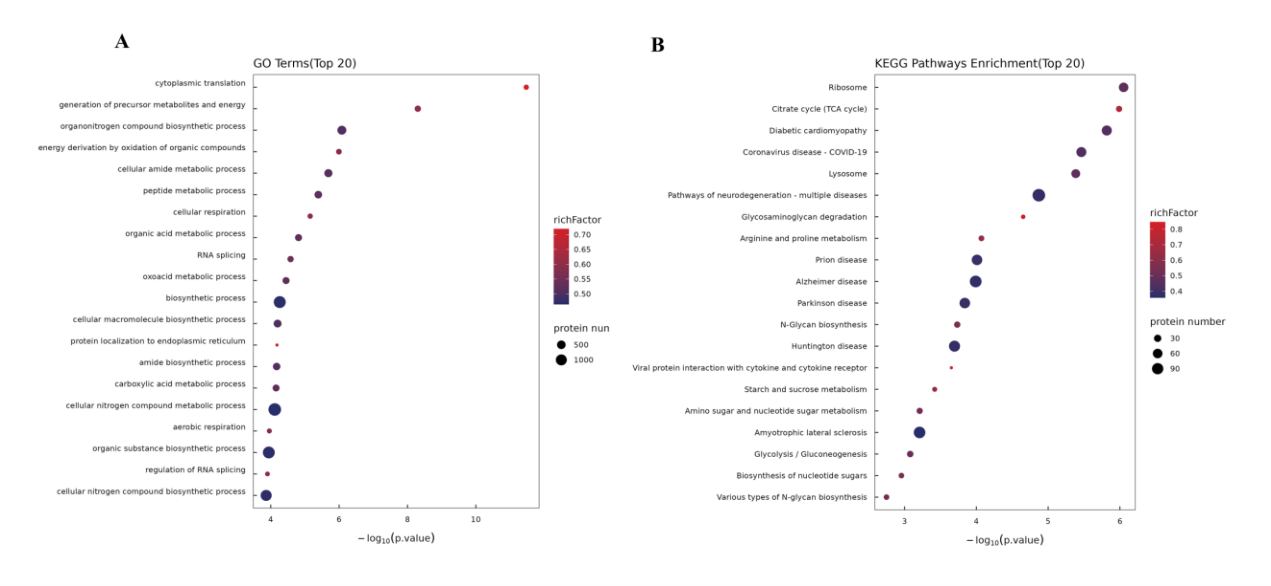 |
| --- | --- |

**Supplemental Figure 8. Functional annotation of *GTDC1* knockout-induced proteome dysregulation.**

(A) GO enrichment for biological processes among differentially abundant proteins in *GTDC1*^‾/‾^ vs. WT 2BS cells. Bubble size indicates protein count; color scale reflects statistical significance (-log_10_[FDR]).

(B) KEGG pathway enrichment of dysregulated proteins.

**Supplemental Table 1**

UHPLC-QQQ-MS instrument parameters of plasma metabolites

| **Compound Name** | **Precursor Ion** | **Product Ion** | **Dwell** | **Fragmentor** | **Collision Energy** |
| --- | --- | --- | --- | --- | --- |
| Glucosamine | 180.1 | 84.1 | 30 | 380 | 20 |
| D-glucosamine 1-phosphate | 260.1 | 180.1 | 30 | 380 | 15 |
| 6-Keto-prostaglandin F1α | 371.2 | 273.2 | 30 | 380 | 15 |
| 5-Hydroxy-6,8,11,14,17-eicosapentaenoic acid | 319.1 | 263.1 | 30 | 380 | 15 |
| 11-Hydroxy-5Z,8Z,12E,14Z,17Z-eicosapentaenoic acid | 319.2 | 275.2 | 30 | 380 | 15 |
| ω-Muricholic acid | 409.3 | 345.3 | 30 | 380 | 25 |
| Glutaric acid | 133.0 | 87.0 | 30 | 380 | 15 |
| 2-Methylcitric acid | 207.0 | 143.0 | 30 | 380 | 15 |
| Nicotinamide ribotide | 256.1 | 136.1 | 30 | 380 | 25 |
| Acetylcysteine | 164.0 | 123.0 | 30 | 380 | 15 |
| Petroselinic acid | 283.3 | 97.1 | 30 | 380 | 20 |
| 11Z-Eicosenoic acid | 311.3 | 185.1 | 30 | 380 | 20 |
| 13,14-Dihydro-15-keto Prostaglandin F1α | 353.2 | 309.2 | 30 | 380 | 15 |
| Salicyluric acid | 196.1 | 120.1 | 30 | 380 | 15 |
| Quinoline | 130.1 | 78.0 | 30 | 380 | 15 |
| 2'-Deoxyadenosine 5'-monophosphate | 332.1 | 136.1 | 30 | 380 | 25 |
| Prostaglandin F2b | 355.2 | 337.2 | 30 | 380 | 15 |
| Hyodeoxycholic acid | 393.1 | 161.1 | 30 | 380 | 25 |
| 4,5-Dihydroorotic acid | 159.0 | 113.0 | 30 | 380 | 15 |
| Fumaric acid | 117.0 | 73.0 | 30 | 380 | 15 |

**Supplemental Table 2**

Detailed mass spectrometry information of 52 tRNA fragments from rats using UHPLC-QTOF-MS

| **tRNA** | **RNase T1 digestion product** | **Rt(min)** | **Formula** | **ion type** | **m/z** | **diff(ppm)** | **Deconvoluted mass** |
| --- | --- | --- | --- | --- | --- | --- | --- |
| tRNA^Asn (AAC)^ | AAAG | 6.16 | C_40_O_26_N_20_H_50_P_4_ | [M-H]^-^ | 1349.2100 | -2.02 | 1350.22 |
|  | CCCACCCAG | 8.95 | C_106_O_78_N_36_H_136_P_10_ | [M-3H]^-3^ | 951.1165 | 9.93 | 2581.38 |
|  | CUQUU[t^6^A]ACCG | 10.09 | C_106_O_78_N_36_H_136_P_10_ | [M-5H]^-5^  [M-4H]^-4^  [M-3H]^-3^ [M-2H]^-2^ | 693.2992 866.8747 1156.1678 1734.7537 | -1.19 | 3470.52 |
|  | D[acp^3^U]AG | 2.95 | C_42_O_32_N_15_H_57_P_4_ | [M-H]^-^ | 1406.2175 | 0.49 | 1407.22 |
|  | UCUCUG | 6.48 | C_55_O_46_N_17_H_71_P_6_ | [M-3H]^-3^  [M-2H]^-2^  [M-H]^-^ | 629.4000 944.6100 1890.2300 | -6.42 | 1891.23 |
| tRNA^Asn (AAU)^ | UU[t^6^A]ACCG | 8.51 | C_71_O_54_N_26_H_91_P_7_ | [M-3H]^-3^ | 795.1052 | 2.09 | 2388.32 |
| tRNA^Asp (GAC)^ | A[m^5^C][m^5^C]G | 4.77 | C_40_O_28_N_16_H_54_P_4_ | [M-2H]^-2^ [M-H]^-^ | 664.1000 1329.2100 | 6.29 | 1330.22 |
|  | AUUCCCCG | 8.45 | C_74_O_58_N_26_H_96_P_8_ | [M-4H]^-4^  [M-3H]^-3^ [M-2H]^-2^ | 630.0800 840.4400 1261.1600 | -0.14 | 2524.33 |
|  | CUCG | 5.95 | C_37_O_30_N_13_H_49_P_4_ | [M-2H]^-2^ [M-H]^-^ | 638.5700 1278.1500 | 4.59 | 1279.16 |
|  | TΨCG | 5.6 | C_38_O_31_N_12_H_50_P_4_ | [M-2H]^-2^ [M-H]^-^ | 646.0800 1293.1600 | -1.61 | 1294.17 |
|  | UAΨAG | 5.11 | C_48_O_36_N_19_H_60_P_5_ | [M-2H]^-2^ [M-H]^-^ | 815.6012 1632.2056 | -6.96 | 1633.22 |
| tRNA^Asp (GAC)^ | UCA[m^5^C]G | 3.54 | C_48_O_36_N_18_H_63_P_5_ | [M-2H]^-2^  [M-H]^-^ | 810.1100 1621.2300 | -0.37 | 1622.23 |
| tRNA^Asp (manQAC)^ | CUC[manQ]UCA[m^5^C]G | 7.83 | C_98_O_72_N_31_H_130_P_9_ | [M-5H]^-5^ [M-4H]^-4^ [M-3H]^-3^  [M-2H]^-2^ | 633.4990 792.1230 1056.4989 1585.2505 | -1.8 | 3171.52 |
| tRNA^Gln (CAA)^ | AAΨCCAG | 8.09 | C_67_O_48_N_28_H_85_P_7_ | [M-3H]^-3^ | 754.4357 | -1.02 | 2266.32 |
|  | A[Cm]UCUG | 7.77 | C_65_O_51_N_23_H_84_P_7_ | [M-2H]^-2^ | 963.1250 | -1.54 | 1928.26 |
|  | CACUCUG | 7.53 | C_65_O_51_N_23_H_84_P_7_ | [M-3H]^-3^  [M-2H]^-2^ | 739.0900 1109.1400 | 0.31 | 2219.28 |
|  | DDAG | 6.35 | C_38_O_30_N_14_H_52_P_4_ | [M-2H]^-2^  [M-H]^-^ | 653.0900 1307.1800 | -0.85 | 1308.19 |
|  | ΨΨCA[m^1^A]AUCUCG | 9.59 | C_104_O_79_N_37_H_132_P_11_ | [M-5H]^-5^  [M-4H]^-4^  [M-3H]^-3^ [M-2H]^-2^ | 699.6900 874.8600 1167.1500 1751.2300 | -5.17 | 3503.47 |
| tRNA^Glu (GAA)^ | [m^5^Um]ΨCG | 6.35 | C_39_O_31_N_12_H_52_P_4_ | [M-2H]^-2^  [M-H]^-^ | 653.0900 1307.1800 | -4.24 | 1308.19 |
|  | ACUCCCG | 7.53 | C_65_O_50_N_24_H_85_P_7_ | [M-2H]^-2^  [M-H]^-^ | 803.5985 1608.2005 | 7.43 | 2218.28 |
|  | AUUCCUG | 7.53 | C_65_O_52_N_22_H_83_P_7_ | [M-3H]^-3^  [M-2H]^-2^ | 739.0900 1109.1300 | -4.39 | 2220.28 |
|  | ΨU[mcm^5^s^2^U]UCACCCAG | 11.26 | C_105_O_81_N_35_H_134_S_1_P_11_ | [M-5H]^-5^  [M-4H]^-4^  [M-3H]^-3^ [M-2H]^-2^ | 709.8800 887.6000 1183.8000 1776.2100 | -7.55 | 3553.45 |
| tRNA^Glu (GAA)^ | UCΨAG | 6.27 | C_47_O_37_N_17_H_60_P_5_ | [M-2H]^-2^  [M-H]^-^ | 803.6000 1608.2000 | -5.7 | 1609.21 |
| tRNA^Leu (CAU)^ | [m^1^A]AUCCCACUUCUG | 11.33 | C_122_O_93_N_43_H_156_P_13_ | [M-2H]^-2^  [M-H]^-^ | 650.5798 1302.1650 | -4.11 | 4113.55 |
|  | ACUAAΨΨCUG | 4.63 | C_94_O_72_N_34_H_118_P_10_ | [M-5H]^-5^  [M-2H]^-2^ | 636.0793 1591.1971 | -4.95 | 3184.41 |
|  | DCΨAAG | 6.48 | C_57_O_43_N_22_H_74_P_6_ | [M-3H]^-3^  [M-2H]^-2^ [M-H]^-^ | 645.7500 969.1300 1939.2600 | -3.56 | 1940.28 |
|  | UCAG | 6.27 | C_38_O_29_N_15_H_49_P_4_ | [M-H]^-^ | 1302.1700 | -0.08 | 1303.18 |
| tRNA^Lys (AAA)^ | CCCCACG | 7.39 | C_65_O_49_N_25_H_86_P_7_ | [M-3H]^-3^-H_2_O | 732.0934 | 3.26 | 2217.30 |
| tRNA^Lys (AAG)^ | [m^5^Um]ΨCA[m^1^A]G | 6.91 | C_60_O_43_N_22_H_78_P_6_ | [M-3H]^-3^  [M-2H]^-2^ [M-H]^-^ | 659.1000 990.1400 1981.2700 | 5.59 | 1980.29 |
|  | ACU[mcm^5^s^2^U]UU[ms^2^t^6^A]AΨCUG | 13.06 | C_121_O_93_N_39_H_153_S_2_P_12_ | [M-6H]^-6^  [M-5H]^-5^  [M-4H]^-4^  [M-3H]^-3^ | 678.5800 814.2900 1018.3700 1357.8200 | -3.71 | 4075.49 |
|  | CAΨCAG | 6.82 | C_57_O_42_N_23_H_73_P_6_ | [M-3H]^-3^  [M-H]^-^ | 644.7500 1936.2600 | -2.81 | 1937.28 |
|  | UUCG | 2.9 | C_37_O_31_N_12_H_48_P_4_ | [M-2H]^-2^  [M-H]^-^ | 639.0700 1279.1500 | -6.23 | 1280.16 |
| tRNA^Ser (AGU)^ | A[i^6^A]A[Ψm]CCAU[Um]G | 17.6 | C_102_O_70_N_37_H_131_P_10_ | [M-5H]^-5^  [M-4H]^-4^  [M-3H]^-3^ [M-2H]^-2^ | 659.9000 825.1300 1100.5000 1651.2600 | -3.04 | 3303.53 |
| tRNA^Ser (AGU)^ | A[m^3^C]UIG | 6.82 | C_49_O_36_N_19_H_62_P_5_ | [M-2H]^-2^  [M-H]^-^ | 822.6107 1646.2243 | -5.37 | 1647.23 |
|  | C[m^2^_2_G]AΨG | 6.44 | C_50_O_36_N_20_H_65_P_5_ | [M-H]^-^ | 1676.2100 | -0.75 | 1676.26 |
|  | DDAAG | 6.22 | C_48_O_36_N_19_H_64_P_5_ | [M-2H]^-2^  [M-H]^-^ | 817.6200 1636.2300 | 1.96 | 1637.24 |
|  | U[m^3^C]UCCCCG | 7.91 | C_74_O_59_N_24_H_98_P_8_ | [M-4H]^-4^ | 927.8300 | -1.42 | 2514.33 |
|  | [m^1^A]AUCCCAUCCUCG | 11.39 | C_122_O_92_N_44_H_157_P_13_ | [M-6H]^-6^  [M-5H]^-5^  [M-4H]^-4^  [M-3H]^-3^ | 684.7500 821.9100 1027.6300 1370.5100 | -1.39 | 4112.56 |
|  | A[m^3^C]UG | 5.6 | C_39_O_29_N_15_H_51_P_4_ | [M-2H]^-2^  [M-H]^-^ | 657.5900 1316.1800 | 6.39 | 1317.18 |
|  | CU[m^3^C]UG | 4.47 | C_47_O_38_N_15_H_62_P_5_ | [M-2H]^-2^  [M-H]^-^ | 798.6000 1598.2000 | -3.51 | 1599.21 |
|  | CU[m^6^t^6^A]AΨCCAU[Um]G | 11.77 | C_110_O_83_N_38_H_141_P_11_ | [M-5H]^-5^  [M-4H]^-4^  [M-3H]^-3^ [M-2H]^-2^ | 731.7000 914.8700 1220.1700 1830.7500 | -2.06 | 3662.52 |
| tRNA^Val (GUU)^ | [m^1^A]AACCG | 6.52 | C_59_O_40_N_26_H_76_P_6_ | [M-3H]^-3^  [M-2H]^-2^ [M-H]^-^ | 657.1000 986.1500 1973.2900 | 4.01 | 1974.31 |
|  | DDAUCAC[m^2^G]ΨUCG | 17.53 | C_113_O_88_N_39_H_147_P_12_ | [M-4H]^-4^  [M-3H]^-3^ [M-2H]^-2^ | 956.8700 1276.5000 1914.7400 | -0.14 | 3829.51 |
| mit-tRNA^Asp (GAC)^ | UUAAG | 5.09 | C_48_O_36_N_19_H_60_P_5_ | [M-2H]^-2^  [M-H]^-^ | 815.6012 1632.2056 | -6.96 | 1633.22 |
| mit-tRNA^Arg (AGA)^ | ACΨCAUUAG | 9.7 | C_85_O_64_N_32_H_107_P_9_ | [M-4H]^-4^  [M-3H]^-3^ [M-2H]^-2^ | 718.8400 958.7900 1438.6800 | -0.07 | 2878.37 |
| mit-tRNA^Leu (CAU)^ | CAACUCCAAAUAAAAG | 13.24 | C_154_O_106_N_66_H_192_P_16_ | [M-7H]^-7^  [M-5H]^-5^  [M-4H]^-4^  [M-3H]^-3^ | 735.9617 1030.5481 1288.4304 1718.5776 | -0.02 | 5156.74 |
| mit-tRNA^Phe (UUU)^ | AA[ms^2^i^6^A]AΨG | 19.52 | C_65_O_40_N_27_H_83_S_1_P_6_ | [M-3H]^-3^ [M-2H]^-2^ | 698.7800 1048.6700 | -2.13 | 2099.35 |
|  | CAAAG | 6.36 | C_49_O_33_N_23_H_62_P_5_ | [M-2H]^-2^  [M-H]^-^ | 827.1200 1657.2500 | 4.55 | 1655.25 |
|  | CUUAG | 5.01 | C_47_O_37_N_17_H_60_P_5_ | [M-2H]^-2^  [M-H]^-^ | 803.6000 1608.2000 | -5.7 | 1609.21 |
| mit-tRNA^Trp (UGG)^ | AUAUACAG | 9.33 | C_77_O_55_N_32_H_96_P_8_ | [M-4H]^-4^  [M-3H]^-3^ [M-2H]^-2^ | 648.0900 864.7800 1297.6700 | 0.3 | 2596.36 |
|  | CCCUUAG | 7.53 | C_65_O_51_N_23_H_84_P_7_ | [M-3H]^-3^ [M-2H]^-2^ | 739.0900 1109.1400 | 0.31 | 2219.28 |
| mit-tRNA^Val (GUU)^ | CAΨCUG | 6.52 | C_56_O_44_N_20_H_72_P_6_ | [M-3H]^-3^  [M-2H]^-2^ [M-H]^-^ | 637.4100 956.1200 1914.2400 | -2.1 | 1914.25 |
